# Probing the SAM Binding Site of SARS-CoV-2 nsp14 in vitro Using SAM Competitive Inhibitors Guides Developing Selective bi-substrate Inhibitors

**DOI:** 10.1101/2021.02.19.424337

**Authors:** Kanchan Devkota, Matthieu Schapira, Sumera Perveen, Aliakbar Khalili Yazdi, Fengling Li, Irene Chau, Pegah Ghiabi, Taraneh Hajian, Peter Loppnau, Albina Bolotokova, Karla J.F. Satchell, Ke Wang, Deyao Li, Jing Liu, David Smil, Minkui Luo, Jian Jin, Paul V. Fish, Peter J. Brown, Masoud Vedadi

## Abstract

The COVID-19 pandemic has clearly brought the healthcare systems world-wide to a breaking point along with devastating socioeconomic consequences. The SARS-CoV-2 virus which causes the disease uses RNA capping to evade the human immune system. Non-structural protein (nsp) 14 is one of the 16 nsps in SARS-CoV-2 and catalyzes the methylation of the viral RNA at N7-guanosine in the cap formation process. To discover small molecule inhibitors of nsp14 methyltransferase (MT) activity, we developed and employed a radiometric MT assay to screen a library of 161 in house synthesized S-adenosylmethionine (SAM) competitive methyltransferase inhibitors and SAM analogs. Among seven identified screening hits, SS148 inhibited nsp14 MT activity with an IC_50_ value of 70 ± 6 nM and was selective against 20 human protein lysine methyltransferases indicating significant differences in SAM binding sites. Interestingly, DS0464 with IC_50_ value of 1.1 ± 0.2 μM showed a bi-substrate competitive inhibitor mechanism of action. Modeling the binding of this compound to nsp14 suggests that the terminal phenyl group extends into the RNA binding site. DS0464 was also selective against 28 out of 33 RNA, DNA, and protein methyltransferases. The structure-activity relationship provided by these compounds should guide the optimization of selective bi-substrate nsp14 inhibitors and may provide a path towards a novel class of antivirals against COVID-19, and possibly other coronaviruses.

## Introduction

COVID-19, a severe acute respiratory syndrome in humans, is caused by SARS-CoV-2. It first surfaced in December 2019^1^, and soon became a pandemic with more than 70 million confirmed cases and more than 1.5 million deaths reported world-wide to-date (https://www.who.int/emergencies/diseases/novel-coronavirus-2019). SARS-CoV-2 belongs to the *Coronaviridae* family of viruses^2,3^ that also includes the Severe Acute Respiratory Syndrome Coronavirus (SARS-CoV) and Middle Eastern Respiratory Syndrome Coronavirus (MERS-CoV) which caused SARS and MERS epidemics in 2002 and 2012, respectively.^4,5^ SARS-CoV-2 contains a non-segmented, positive sense 30 kb RNA that consists of 14 open reading frames (ORFs)^6^, encoding 16 non-structural proteins (nsp) and four main structural and accessory proteins.^7^ The 16 non-structural proteins (referred to as nsp1 to nsp16) are more conserved amongst coronaviruses compared to the structural and accessory proteins.^8^ These nsps in coronaviruses form a replicase-transcriptase complex and are essential for the transcription and replication of the virus.^9^ Among these, nsp14 and nsp16 are RNA methyltransferases involved in RNA capping.^10^

Nsp14 is a bi-functional protein with a C-terminal methyltransferase domain catalyzing N7-guanosine methylation, and an N-terminal exoribonuclease domain (**Supplementary Fig. 1**). In the replicase-transcriptase complex of coronaviruses, nsp14 functions as an exoribonuclease and is involved in maintaining the fidelity of coronavirus RNA synthesis.^11^ Nsp14 in complex with nsp10 can function as a proofreading exoribonuclease and removes 3’-end mismatched nucleotides from dsRNA.^12^ Breaking this interaction between nsp10 and nsp14 results in a decrease in virus replication fidelity.^13^ Besides nsp10, nsp14 also interacts with the nsp7-nsp8-nsp12 complex where the exonuclease function of nsp14 decreases the incidence of mismatched nucleotides^14^ by erasing the mutated nucleotides.^11^

While complex formation between nsp10 and nsp14 is required for enhanced exoribonuclease activity, the MT activity of nsp14 is independent of nsp10-nsp14 complex formation.^15,16^ The nsp14 SAM-dependent methyltransferase (MTase) activity is essential for viral mRNA capping.^16,17^ The cap1 structure at the 5’-end of viral RNA helps in masking the virus from the host immune system.^18,19^ Cap (GpppN) structure in nascent RNA of coronaviruses is formed by nsp13^20,21^ and a guanylyltransferase (GTase). Nsp14 methylates this cap structure at the N7 position of the guanosine, forming a cap-O (N7mGpppN).^17^ Nsp16 further 2’-O-methylates the product of the nsp14 methyltransferase activity, completing the capping process (N7mGpppNm).^16,22^

Nsp14 is conserved amongst the seven coronaviruses known to infect humans to-date (**Supplementary Fig. 2**).^23^ The SARS-CoV-2 nsp14 overall amino acid sequence shows 95.1, 62.7, 57.8, 58.5, 52.9 and 53.7% identity with nsp14 from SARS-CoV, MERS-CoV, OC43, HKU1, 299E and NL63, respectively. This suggests the possibility of SARS-CoV-2 nsp14 inhibitors also inhibiting nsp14 methyltransferase activity of other coronaviruses. Such pan inhibitors would be priceless for developing pan anti-viral therapeutics for COVID-19 that would be also effective on future coronaviruses which may jump to humans. In this study, we first developed a radiometric high throughput activity assay for SARS-CoV-2 nsp14 methyltransferase activity and screened a library of 161 S-adenosylmethionine (SAM) competitive methyltransferase inhibitors we previously synthesized, and SAM analogs. We identified seven reproducible hits which we mapped on the active site of SARS-CoV-2 nsp14. The data shows for the first time a clear path towards the development of potent and selective bi-substrate nsp14 inhibitors that may lead to a novel class of therapeutics for COVID-19 and possibly other coronaviruses to come.

## Results

### Assay development and optimization for high throughput screening

Developing therapeutics for COVID-19 and other coronaviruses requires reliable high throughput screening assays. Radiometric assays have been widely used for developing potent substrate and SAM competitive inhibitors for human methyltransferases within the last decade.^24^ Using biotinylated RNA substrate, a radiometric nsp14 methyltransferase (MTase) activity assay was developed with ^3^H-SAM as a methyl donor (**Fig. 1**). Nsp14 MTase activity was evaluated in various buffers, pH and additives (**Supplementary Fig. 3**). The highest MTase activity was observed in Tris HCl buffer at pH 7.5. No significant effect was observed for DTT up to 10 mM and it was included in the assay reaction mixture at 5 mM to maintain reducing conditions. Triton X-100 at 0.01% was added to the reaction buffer to minimize binding of proteins and compounds to plates. DMSO at concentrations up to 10% had little effect on nsp14 MTase activity. However, MgCl2 was only tolerated at concentrations below 1 mM (**Supplementary Fig. 3**). The assay optimization resulted in selecting 20 mM Tris HCl pH 7.5, 250 μM MgCl2, 5 mM DTT and 0.01 % Triton X-100 for testing nsp14 MTase activity and determining its kinetic parameters.

**Figure 1:**
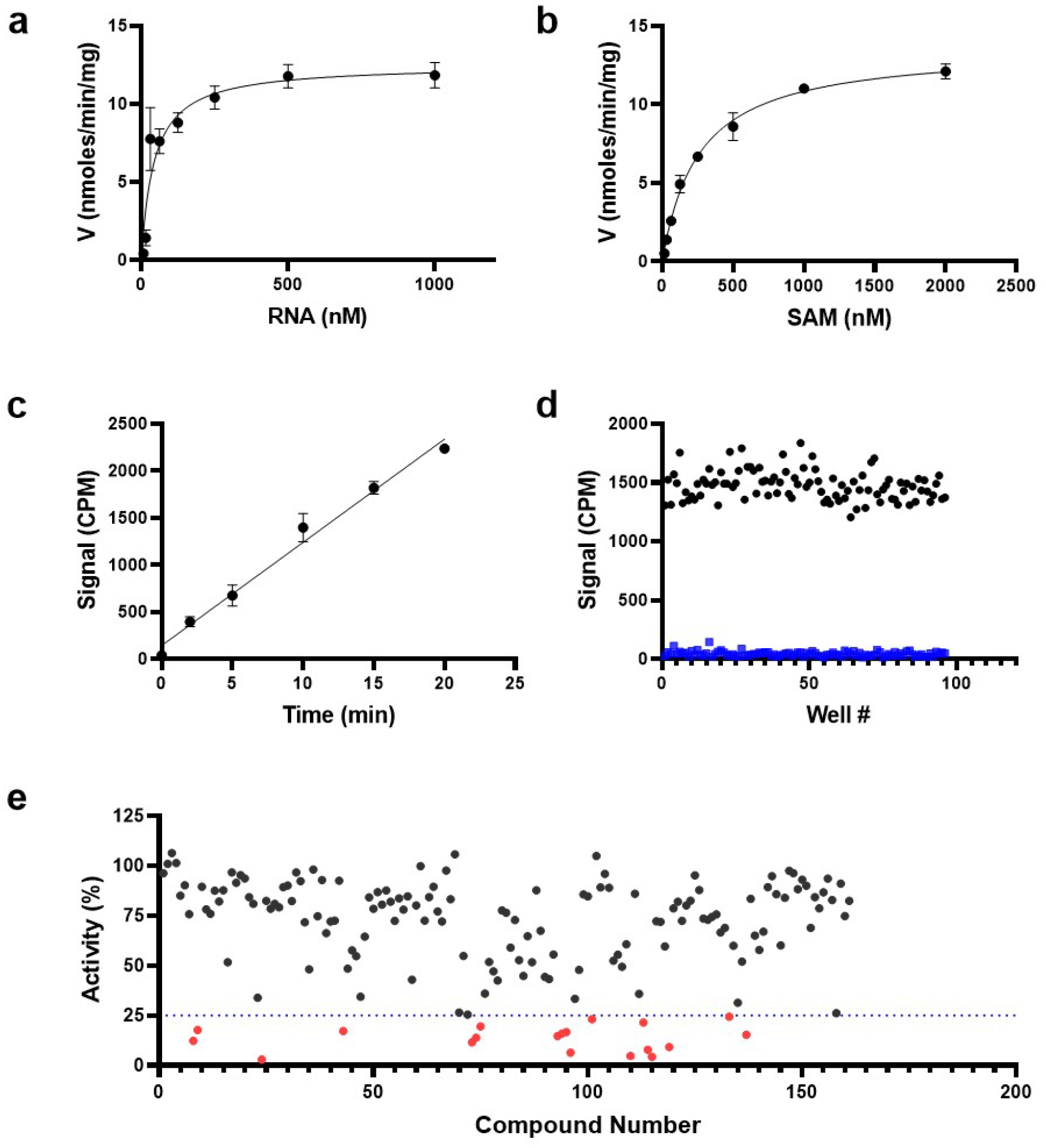
Kinetic characterization and screening of nsp14 methyltransferase activity. The optimized radiometric methyltransferase assay was used to determine the K_m_^app^ and k_cat_^app^ values for (a) RNA and (b) SAM at saturating concentrations of SAM and RNA, respectively. (c) Linearity of initial velocity of nsp14 methyltransferase reaction was tested at K_m_ of both substrates. (d) Z’-factor was determined at the same conditions as in “c”. Nsp14 activity was assessed in the (•) absence (100% activity) and 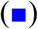 presence of 1 μM Sinefungin. (e) Nsp14 was screened against a library of SAM analogs. A collection of 161 SAM competitive inhibitors and analogs was tested at 50 μM for inhibition of nsp14 activity under conditions described for Z’-factor determination. Compounds with 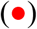 higher than 75% inhibition were selected for follow up experiments. Values presented in a, b, and c are mean ± standard deviation of three independent experiments (n=3). Values in d and e are from single screening.

### Kinetic parameters of SARS CoV-2 nsp14 MTase activity

Using optimized assay conditions, linearity of initial velocities (activity versus time) was assessed at various concentrations of RNA at fixed SAM concentration (1 μM) (**Supplementary Fig. 4a**), and at various concentrations of SAM at fixed concentration of RNA (1 μM) (**Supplementary Fig. 4b**). The slopes of the linear initial velocities were calculated and were used to determine kinetic parameters for nsp14 MTase activity. Apparent K_m_ (K_m_^app^) values of 43 ± 15 nM and 257 ± 20 nM were determined for RNA and SAM, respectively. The calculated k_cat_^app^ values from the two sets of experiments, 48 ± 4 h^-1^ (RNA; **Fig 1a**) and 52 ± 1 h^-1^ (SAM; **Fig 1b**), were reasonably close. Addition of nsp10 at various molar ratios from 1 (nsp10):1 (nsp14) to 20 (nsp10):1 (nsp14) did not have any significant effect on nsp14 methyltransferase activity (**Supplementary Fig. 5**).

### Assay amenability to high throughput screening

In small molecule screening campaigns, typically the assays are performed at K_m_ of the substrates to allow potential inhibitors to compete with the substrates and allow their binding to be detected. However, the activity of the enzyme should be linear during the assay period. As the radiometric methyltransferase assays are endpoint assays, lack of linearity may mask inhibition of some compounds. Testing the activity of nsp14 at 50 μM RNA and 250 nM SAM indicated that the assay can be run for at least 20 minutes while maintaining the linearity (**Fig. 1c**). To determine the reproducibility of such conditions for high throughput screening, the assay was performed in the presence (1 μM) and absence of sinefungin, a pan methyltransferase inhibitor that inhibits nsp14 MTase activity with an IC_50_ value of 0.019 ± 0.01 μM (**Supplementary Fig. 6**). A Z’-Factor of 0.69 was calculated for nsp14 screening indicating suitability of the assay for high throughput screening (**Fig. 1d**).

### Screening SARS CoV-2 nsp14 against a collection of potential SAM competitive inhibitors

An in-house library of 161 SAM competitive methyltransferase inhibitors and SAM analogs was screened against nsp14 at 50 μM, and 19 compounds were identified that inhibited nsp14 MTase activity more than 75% (**Fig. 1e**). Compounds that interfered with the readout signal or were not reproducible were eliminated. The remaining seven compounds (**Fig. 2**) inhibited nsp14 MTase activity with IC_50_ values ranging from 70 nM to 95 μM (**Table 1, Fig. 3**). We also tested these compounds for binding to nsp14 by Surface Plasmon Resonance (SPR). Binding was detected for all hits and K_D_ values were calculated for all, except WZ16 with its relatively linear fitting which was not reliable for accurate K_D_ calculation (**Table 1, Supplementary Fig. 7**).

**Figure 2:**
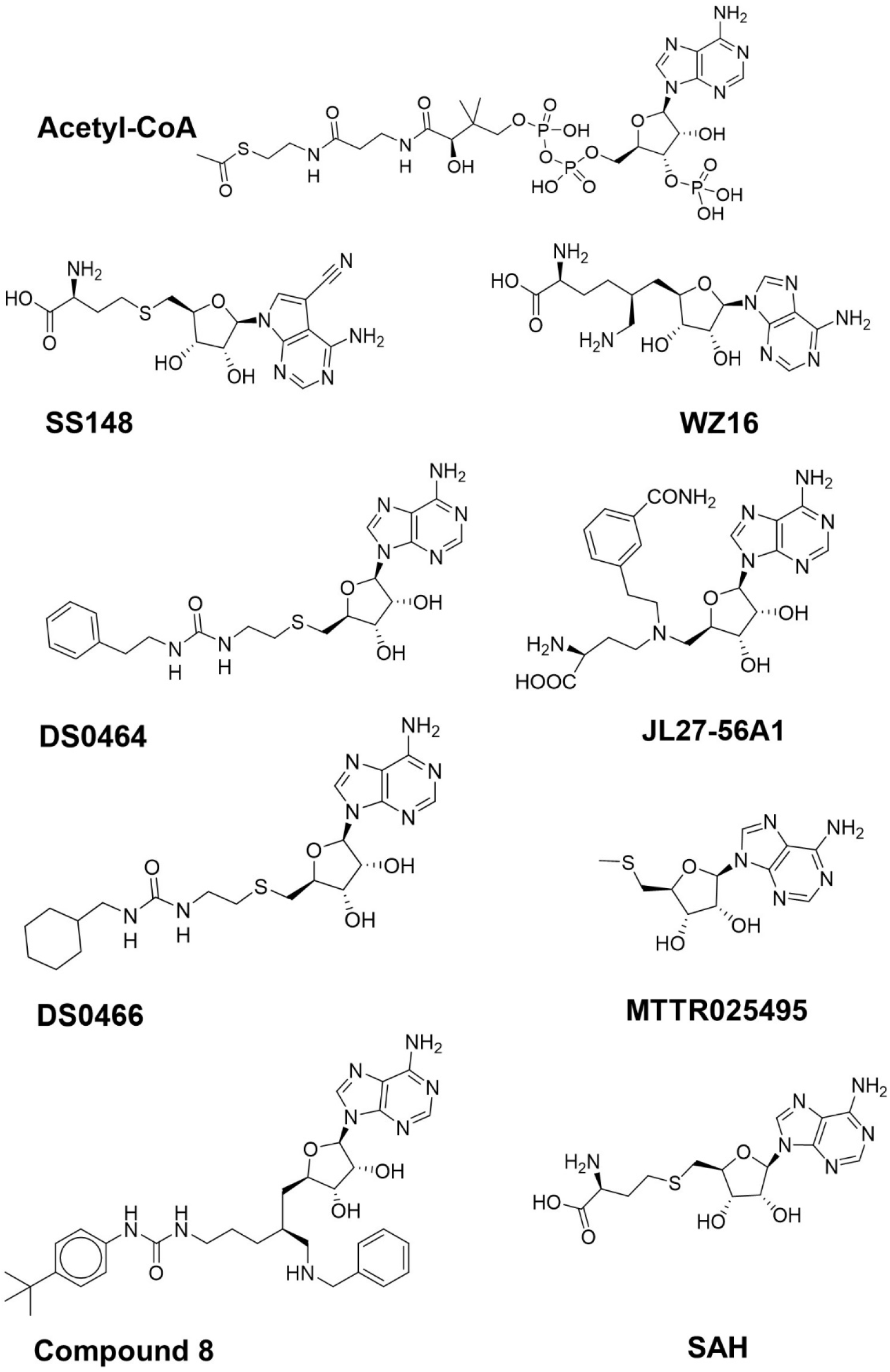
Structures of nsp14 screening hits.

**Figure 3:**
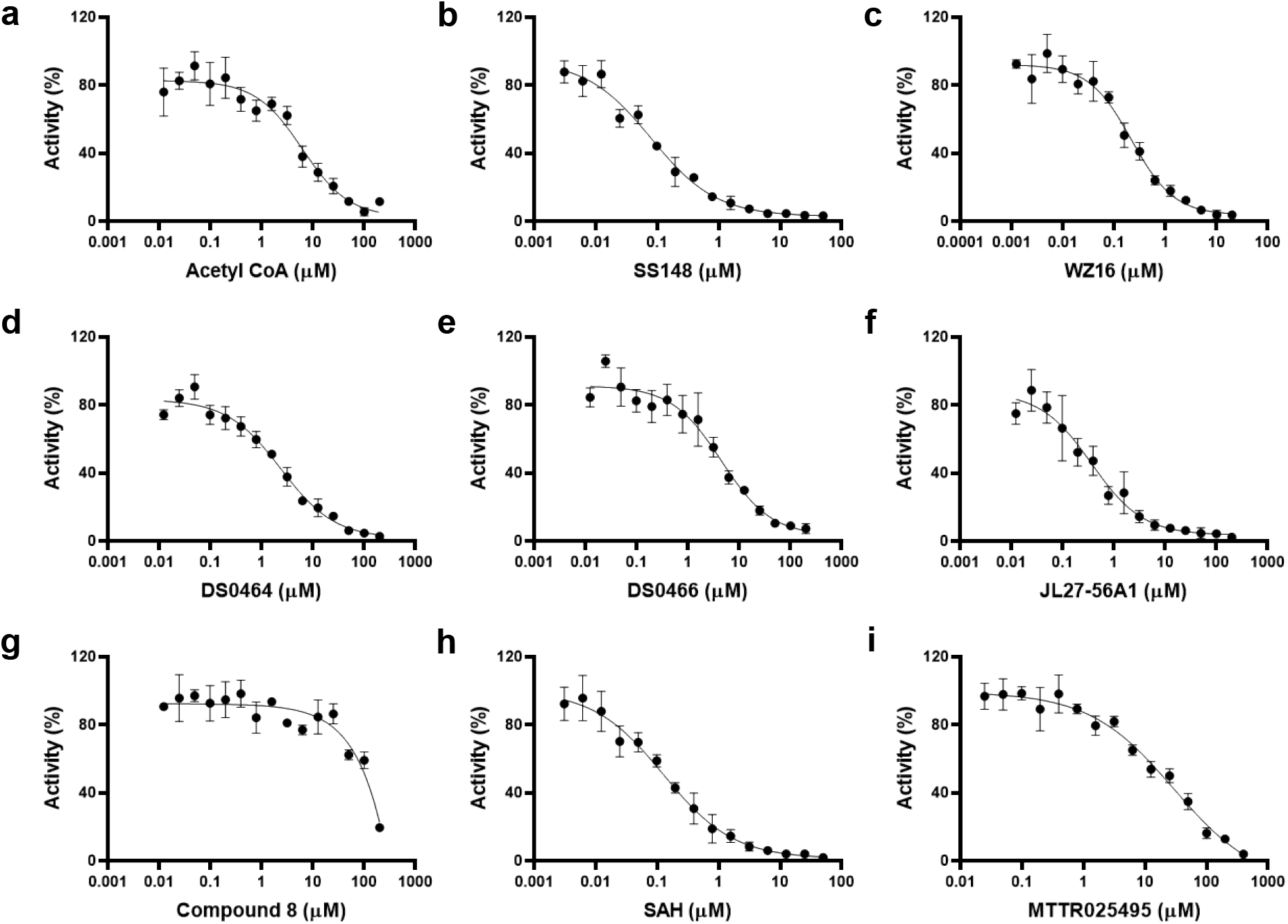
Dose response curves of nsp14 screening hits. The IC_50_ values were determined for (a) acetyl CoA, (b) SS148, (c) WZ16, (d) DS0464, (e) DS0466, (f) JL27-56A1, (g) Compound 8, (h) SAH and (i) MTTR025495 by testing their inhibitory effects ranging from 3 nM to 200 μM final concentration. IC_50_ values are presented in **Table 1**. All values are mean ± standard deviation of three independent experiments (n=3).

**Table 1:** Confirmation and selectivity of nsp14 screening hits. The screening hits were tested for binding to nsp14 by SPR and for inhibition of methyltransferase activity of nsp14 and selected methyltransferases by activity assays. All values are from experiments presented in Figure 3, and Supplementary Fig. 7, 10, 11, 12, 13 and 14.

| Compound | SPR K<sub>D</sub><br>(μM) | IC<sub>50</sub> (μM) | | | | | |
| --- | --- | --- | --- | --- | --- | --- | --- |
|  | nsp14 | nsp 14 | G9a | BCDIN3D | ALKBH5 | SETD3 | METTL3-14 |
| Acetyl CoA | 1.2 | 3 ± 1.3 | NI | NI | NI | NI | 39 |
| SS148 | 0.05 | 0.07 ± 0.006 | NI | 0.03 ± 0.002 | NI | NI | 0.9 |
| WZ16 | NA | 0.19 ± 0.04 | NI | 1.7 | NI | 3.4 | 3.4 |
| DS0464 | 1.6 | 1.1 ± 0.2 | NI | 46 ± 9 | NI | NI | NI |
| DS0466 | 2.4 | 3.4 ± 1.4 | NI | > 50 | NI | NI | NI |
| JL27-56A1 | 0.2 | 0.26 ± 0.13 | NI | 19 | NI | 23 | 10 |
| Compound 8 | 18.9 | 95 ± 6 | NI | NI | NI | NI | NI |
| SAH | 0.14 | 0.13 ± 0.03 | NA | NA | NA | NA | NA |
| MTTR025495 | 6.2 | 17 ± 2 | NA | NA | NA | NA | NA |

**NI: No Inhibition; NA: Not Assessed.**

### Docking and modeling

This limited and focused screening exercise provided important insights on the structural chemistry of SARS CoV-2 nsp14 inhibition (**Table 1**). First, we noted that the de-methylated cofactor S-adenosyl-L-homocysteine (SAH) is a very potent inhibitor of nsp14 (IC_50_: 130 ± 30 nM; K_D_: 140 nM). Decorating the adenine scaffold with a nitrile group at position 7 improved the potency (SS148 IC_50_: 70 ± 6 nM, K_D_: 50 nM) while installing an ethylamine in place of the sulfur group of SAH had little effect on potency (WZ16: IC_50_: 190 ± 40 nM). Deleting the amino-acid end of SAH resulted in a 130-fold loss in SARS-CoV-2 nsp14 inhibition (MTTR025495 IC_50_: 17 ± 2 μM, **Table 1, Fig 3**). Interestingly, compound DS0464, where the amino-acid moiety is replaced with a physico-chemically more favorable phenyl-ethyl-urea, retains significant inhibitory activity (IC_50_: 1.1 ± 0.2 μM). Since all residues lining SAH and the substrate RNA cap GpppA in the SARS-CoV-1 nsp14 structure are conserved in SARS-CoV-2 (**Supplementary Fig. 8**), the SARS-CoV-1 structure was used to dock DS0464 (**Fig. 4**). The model revealed a possible arrangement where the adenosine end of the inhibitor overlays with the adenosine of the bound cofactor, and the terminal phenyl group recapitulates stacking interactions observed between the guanine ring of the RNA cap and surrounding residues (Y420, F426, F506, N386) (**Fig. 4**). Such a binding mode suggests a mechanism of action where DS0464 behaves as a bi-substrate inhibitor. This possibility was further tested by performing mechanism of action (MOA) studies with DS0464 which revealed that DS0464 competes against both SAM and RNA and can act as a bi-functional inhibitor (**Fig. 5, Supplementary Fig. 9**).

**Figure 4:**
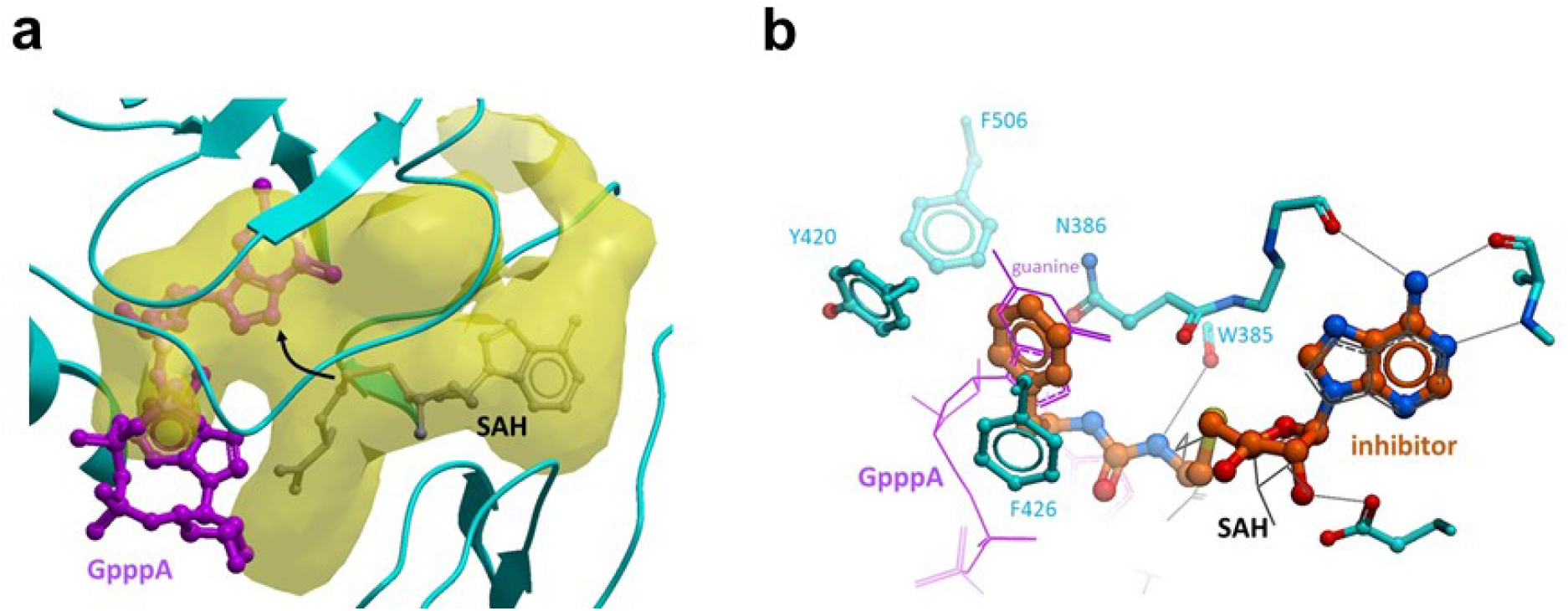
Docking model of DS0464. (a) A large and enclosed binding pocket (yellow) in the methyltransferase domain of nsp14 accommodates both the methyl-donating cofactor (grey - SAH is shown instead) and the methyl-accepting RNA cap GpppA (purple) [PDB code 5C8S]. Black arrow: site of methyl transfer. (b) Docked inhibitor DS0464 occupies the cofactor site and extends into the substrate-binding site.

**Figure 5:**
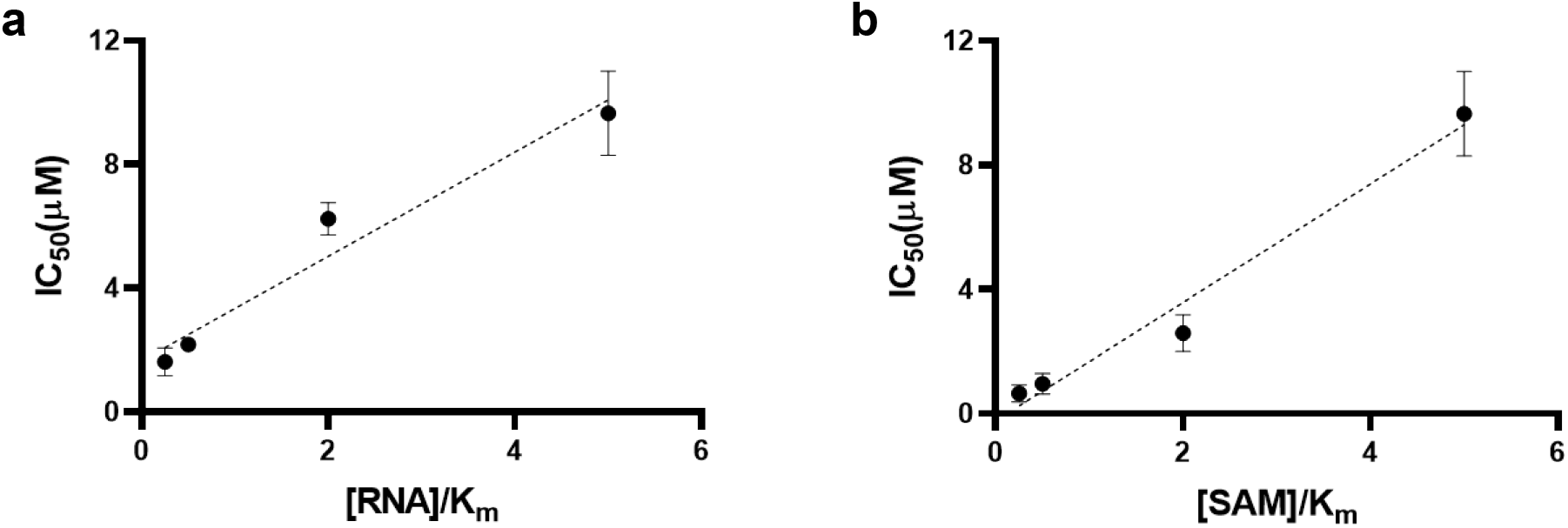
Mechanism of action (MOA) of DS0464. IC_50_ values were determined for DS0464 at (a) varying concentrations of RNA at 1.25 μM of SAM, and (b) varying concentrations of SAM at 250 nM of RNA. DS0464 competes against both RNA and SAM. All values are mean ± standard deviation of three independent experiments (n=3).

### Selectivity of screening hits

The selectivity of all seven compounds was tested against the human RNA methyltransferases BCDIN3D, and METTL3-METTL14 complex (METTL3-14), the RNA demethylase ALKBH5, and protein lysine methyltransferases G9a and SETD3 (**Table 1, Supplementary Fig. 10-14**). Interestingly, none of the compounds inhibited G9a or ALKBH5 activities indicating some level of selectivity. SS148 and DS0464 potently inhibited BCDIN3D with IC_50_ values of 0.03 ± 0.002 and 46 ± 9 μM, respectively, but not G9a, SETD3 or ALKBH5 (**Table 1**). G9a and SETD3 are SET domain methyltransferases that are structurally distinct from class I methyltransferases such as nsp14, BCDIN3D or METTL3-METTL14, which could explain the observed specificity profile.^25^ For instance, the channel separating the substrate and cofactor binding sites is wide in nsp14 but narrow in G9a. Additionally, a cavity that can accommodate the nitrile group of SS148 in nsp14 and BCDIN3D is absent in G9a, in agreement with the obtained IC_50_ values (**Fig 6**). To further characterize our nsp14 inhibitors, the selectivity of SS148 and DS0464 was evaluated against a larger panel of lysine, arginine, DNA and RNA methyltransferases (**Table 2, Fig. 7**). As expected, while SS148 inhibited arginine, DNA and RNA methyltransferases (all class I methyltransferases), it did not inhibit any of the 20 SET domain lysine methyltransferases (**Fig. 7, Table 2, Supplementary Fig. 15**). DS0464 was even more selective and inhibited only PRMT4, PRMT5, PRMT7, DOT1L and BCDIN3D, but none of the protein lysine methyltransferases tested. (**Table 2, Supplementary Fig. 16**).

**Figure 6:**
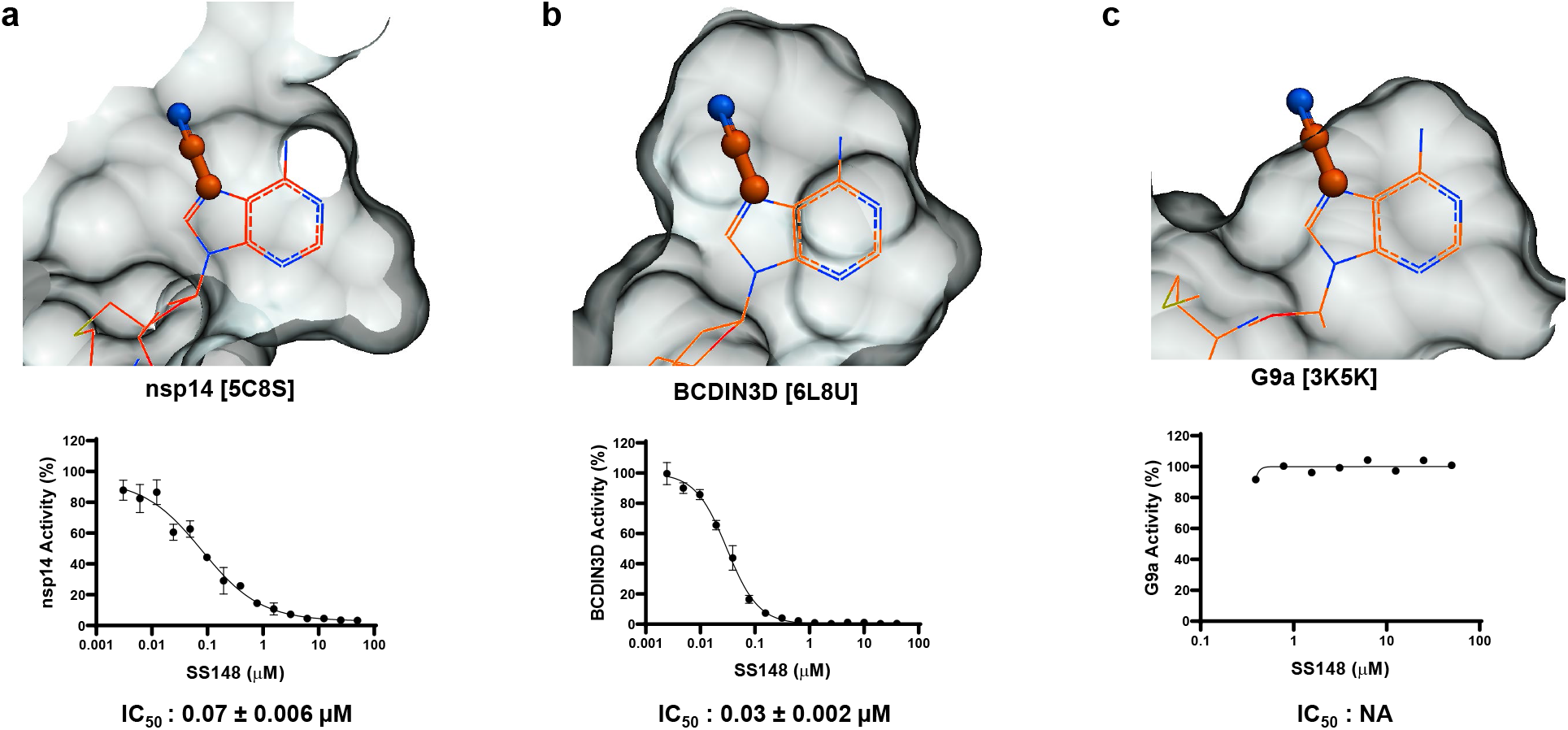
Structural determinant for SS148 selectivity. A nitrile group (stick) at position 7 of the cofactor adenine ring is compatible with the crystal structure of SAH (wire) bound to (a) nsp14 and (b) BCDIN3D, but not (c) G9a. The Van der Waals surface of the cofactor binding pocket is shown as grey mesh. Dose response curves for SS148 against these proteins with their IC_50_ values are also shown.

**Figure 7.**
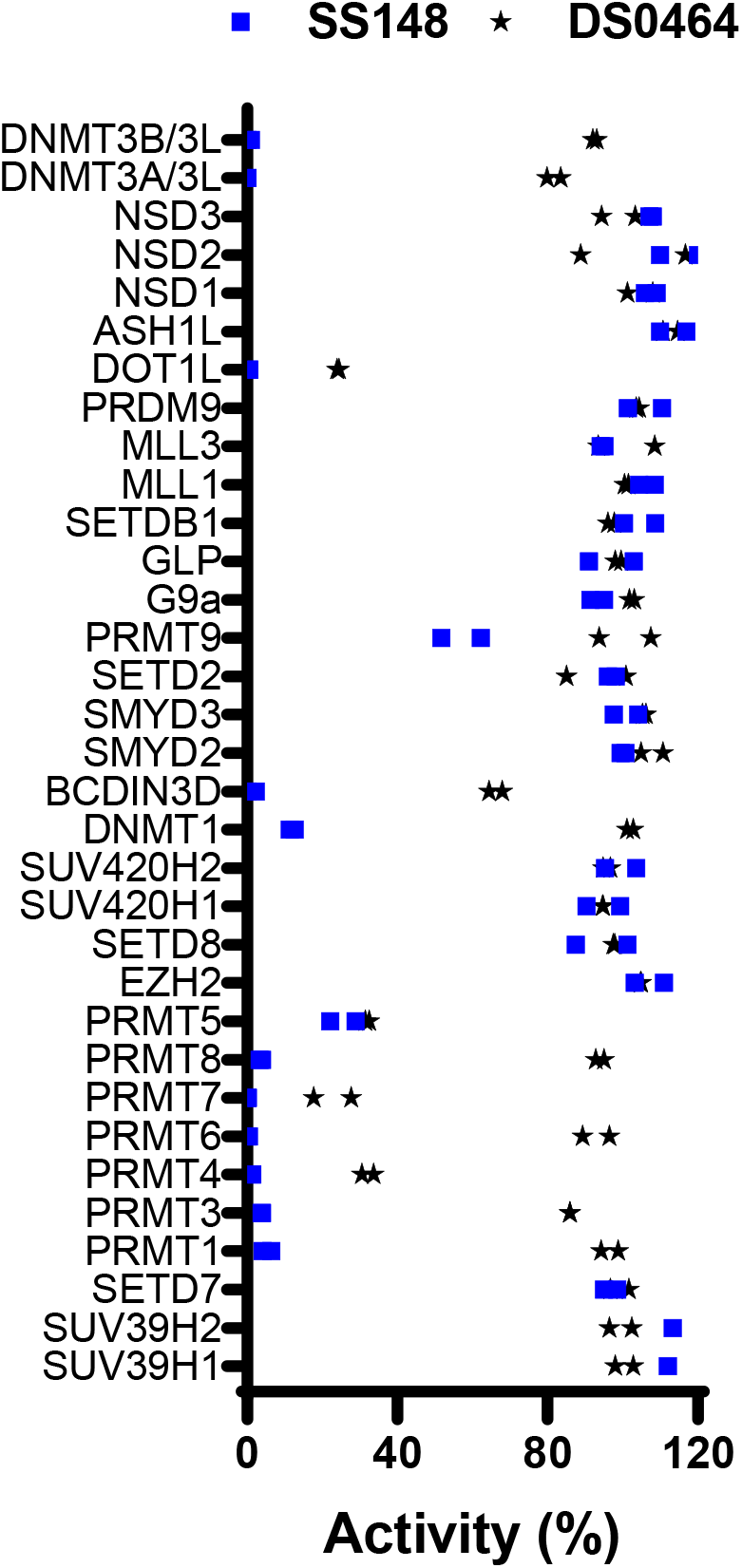
Selectivity of SS148 and DS0464. Both SS148 and DS0464 were screened for selectivity against 33 human RNA, DNA, and protein methyltransferases at 50 μM. All experiments were performed in duplicate. When significant inhibition was observed, IC_50_ values were determined which are presented in **Table 2**.

**Table 2.**
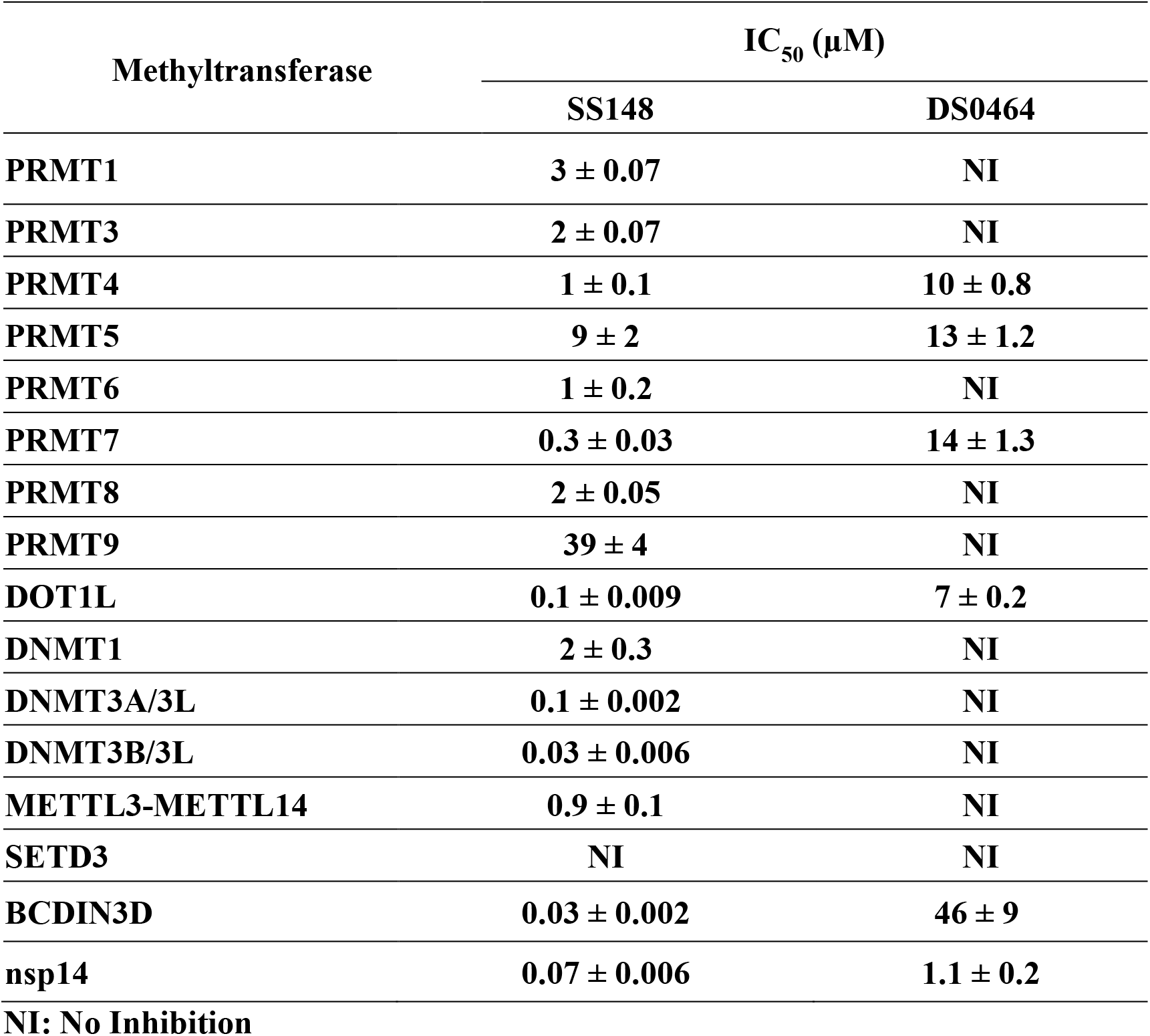
Selectivity of SS148 and DS0464.

## Discussion

The frequent emergence in the last two decades of novel coronaviruses as human pathogens, highlighted by the current COVID-19 pandemic, urgently needs to be addressed, preferably with pan-coronavirus drugs. Nsp14 is an essential methyltransferase in RNA cap formation which is required for protecting viral RNA and proper replication of coronaviruses. Therefore, targeting nsp14 methyltransferase activity would be a viable option towards developing anti-viral therapeutics.^26^ Methyltransferases are druggable.^27^ In the last decade, a significant number of selective and cell-active small molecules (chemical probes) have been discovered for human methyltransferases^24,28,29^ and some are in clinical trials for various cancers.^28,29^

Key to a successful discovery campaign of such chemical probes is the availability of reliable screening methods that could enable medium to high throughput screening with low false-positive and false-negative rates. Various assays including mass spectrometry^30–32^, fluorescence^33^, and radiometric assays^24^ have been used for screening libraries of compounds. Mass spectrometrybased assays require more expensive instrumentation and expertise. Fluorescence assays can be performed in any lab, however, many fluorescent compounds in chemical libraries may increase the background and lead to high numbers of false positives to triage following screening large libraries. Radiometric assays are typically more reliable and have fewer false positives leading to identifying more reliable screening hits.^24^ In this study, we have developed a radiometric assay for nsp14 and employed this assay for screening a small library of selected SAM competitive inhibitors and analogs.

Targeting the SAM binding site has successfully led to the discovery of chemical probes for human methyltransferases such as DOT1L^34^,^35^, EZH2/EZH1^36^, and SMYD2^37^. SS148 (nsp14 IC_50_: 70 ± 6 nM) was reported as a DOT1L inhibitor with a nitrile as a non-traditional replacement for heavy halogen atoms.^38^ This is consistent with the selectivity of SS148 against all other protein lysine methyltransferases (PKMTs) due to narrower active sites that could not fit the added nitrile group (**Fig. 4**). Keeping this substitution in designing future nsp14 inhibitors will provide selectivity against PKMTs. WZ16^39^ and Compound **8** are PRMT4 (CARM1) inhibitors. Although Compound **8** is a relatively weak nsp14 inhibitor (IC_50_: 95 ± 6 μM) it is likely cell-permeable (US20180305391A1). JL27-56A1 is known to modestly inhibit nicotinamide N-methyltransferase (NNMT).^40,41^ Optimization of these series for NNMT resulted in discovery of MS2734, a bi-substrate inhibitor. DS0464 and DS0466 were originally designed for DOT1L inhibition.^34^ DS0464, with replacement of the amino-acid moiety with a physico-chemically more favorable phenyl-ethyl-urea, retains significant nsp14 inhibitory activity (IC_50_: 1.1 ± 0.2 μM). Structural modeling and kinetic data both suggest a mechanism of inhibition consistent with bi-substrate competition. These data reveal a path to developing a series of bi-functional inhibitors to increase the potency of nsp14 SAM competitive inhibitors. Interestingly, DS0464 was selective against 28 out of 33 human RNA, DNA, and protein methyltransferases, indicating a path to developing selective bi-substrate inhibitors towards the discovery of therapeutics for COVID-19 and closely related coronaviruses.

## Conclusion

In this study, we developed a radiometric assay for nsp14 methyltransferase activity, determined the kinetic parameters and optimized the assay for high throughput screening. Through limited screening of SAM competitive inhibitors, we identified seven confirmed hits that we used to probe the active site of nsp14. Our study revealed a path towards developing selective bi-substrate inhibitors for nsp14.

## Methods

### Reagents

S-adenosylhomocysteine (SAH) and sinefungin were purchased from Sigma-Aldrich. S-adenosyl-L-methionine, ^3^H-SAM, and the 384-well FlashPlate^®^ PLUS coated with streptavidin and scintillant (cat. #: SMP410A001PK) were purchased from PerkinElmer. Biotin-labeled single-strand RNA (5’ GpppACCCCCCCCC-Biotin 3’), referred to as RNA substrate here on, was custom synthesized by TriLink BioTechnologies, San Diego, USA. All RNA solutions were prepared by solubilizing in nuclease free water in the presence of RNAseOUT™ recombinant ribonuclease inhibitor (Thermo Fisher, Cat. #. 10777019) at a final concentration of 0.4 U/μL. The D-(+) Biotin, [^3^H(G)], referred to as ^3^H-biotin was purchased from PerkinElmer. The biotinylated single and double-stranded miR-145 substrates for BCDIN3D and METTL3-14 were synthesized by Integrated DNA Technologies (Coralville, IA). All peptide substrates were synthesized by Tufts or GenScript, USA. All other reagents were from Sigma-Aldrich.

### Protein expression and purification

Expression and purification of SARS-CoV-2 nsp14 is provided as supplementary data.

### Radiometric assay development

#### Assay optimization

Methyltransferase activity of nsp14 was measured using a radiometric assay. The transfer of ^3^H-methyl group from ^3^H-SAM to the RNA substrate (5’ GpppACCCCCCCCC-Biotin 3’) was monitored using a scintillation proximity assay (SPA). Unless stated otherwise, all the reactions were carried out in 20 μl final volume and in triplicates at room temperature. For assay optimization, several concentrations of nsp14 (0.5 nM to 1 μM) were tested by mixing 1 μM ^3^H-SAM and 1 μM RNA in 20 μl buffer. The reaction was stopped after 30 minutes by adding 20 μl of 7.5 M guanidium chloride and 20 μl of 20 mM Tris HCl pH 8. The overall assay mixture was then transferred to a 384-well FlashPlate (SPA plate) coated with streptavidin/scintillant. After 3 hours, the amount of methylated RNA formed was quantified using a TopCount^®^ (counts per minute, CPM). To test additives and buffers, nsp14 (1.5 nM) was mixed with 50 nM RNA and 250 nM ^3^H-SAM and the reaction was stopped after 30 minutes. Each buffer (Tris, HEPES and BTP) was tested at 20 mM at pH 7.5. Effect of pH was evaluated in Tris pH 6.5 to pH 9.5. Effect of additives was monitored individually by titrating the reaction mixture with varying concentrations of DTT (from 0.1 mM to 100 mM), MgCl2 (from 0.1 mM to 100 mM), Triton X-100 (from 0.01 % to 10%) and DMSO (from 10 % to 1.25 %) and comparing their relative activity to the samples with no additive.

#### Kinetic characterization

The methyltransferase activity of nsp14 was determined using optimum buffer conditions identified (20 mM Tris HCl pH 7.5, 250 μM MgCl2, 5 mM DTT and 0.01 % Triton X-100). Kinetic parameters (K_m_^app^ and k_cat_^app^) of nsp14 were determined using a series of reactions containing nsp14 at saturating concentration of one substrate (1 μM RNA or 1 μM ^3^H-SAM) and varying concentration of the other (from 15.6 nM to 2000 nM for ^3^H-SAM and from 7.8 nM to 1000 nM for RNA). Reactions were stopped at various time points (2, 5, 10, 15, 20 and 30 minutes). Initial velocities of the reaction were calculated from the linear portion of the reaction curves (**Supplementary Fig. 4**) and were plotted as a function of each substrate concentration (^3^H-SAM and RNA) to determine their K_m_^app^ values using Michaelis-Menten equation. Maximum velocity (Vmax) obtained from the Michaelis-Menten plot was used to calculate the apparent turnover number (k_cat_^app^) in each experiment. To determine the optimum reaction time for screening, linearity of the methylation reaction over time was monitored. The reaction mixture containing 1.5 nM nsp14, 250 nM ^3^H-SAM, 50 nM RNA in 20 mM Tris HCl pH 7.5, 250 μM MgCl2, 5 mM DTT and 0.01 % Triton X-100, were incubated for various durations (0, 2, 5, 10, 15, 20, 30 minutes) and the amount of methylated RNA was quantified over time.

#### IC_50_ determination

Compounds were tested at various concentrations from 12 nM to 200 μM final concentration to determine their half maximal inhibitory concentration (IC_50_) values. Potent compounds (SS148, SAH and WZ16) were tested from 3 nM to 50 μM. Final DMSO concentration was 2%. The final reaction mixture consisted of 1.5 nM nsp14, 250 nM ^3^H-SAM, 50 nM RNA in 20 mM Tris HCl pH 7.5, 250 μM MgCl2, 5 mM DTT and 0.01 % Triton X-100. Reaction time was 20 minutes. Data were fitted to Four Parameter Logistic equation using the GraphPad^®^ Prism 8.

#### Z’-Factor determination

To evaluate the effectiveness of the nsp14 assay for screening purposes, the Z’-factor was determined by running 96 different reactions in the presence or absence of 1 μM sinefungin, a known methyltransferase inhibitor. The final reaction consisted of 1.5 nM nsp14, 250 nM, ^3^H-SAM, 50 nM RNA in 20 mM Tris HCl pH 7.5, 250 μM MgCl2, 5 mM DTT and 0.01 % Triton X-100. Reactions were stopped after 20 minutes. Z’-factor was calculated as described by Zhang et al. ^42^

### Screening

Nsp14 was screened against the in-house library of 161 compounds at 50 μM in 1% DMSO. Compounds with inhibition of more than 75% were selected as screening hits for further analysis. The hits were tested for assay signal quenching at 50 μM. The signal was generated using 0.1 μM ^3^H-biotin in 384-well FlashPlate which are coated with streptavidin/scintillant. Compounds that did not quench the signal were tested by dose response against nsp14 at various concentrations from 12 nM to 200 μM final concentration to determine their half maximal inhibitory concentration (IC_50_) values.

### Mechanism of action

Mechanism of action (MOA) of DS0464 was investigated by determining the IC_50_ values of the compound for nsp14 at saturating concentration of one substrate (250 nM RNA or 1.25 μM ^3^H-SAM) and varying concentration of the other (from 62.5 nM to 1.25 μM for ^3^H-SAM and from 12.5 nM to 250 nM for RNA).

### Selectivity assays

Selectivity assays were performed as previously described.^24^ Briefly, compounds were tested at 50 μM in duplicate using radiometric assays. IC_50_ values were determined for compounds with higher than 50% inhibitory effect, as described above.

### Surface plasmon resonance (SPR)

K_D_ values for initial screening hits for nsp14 was determined by Surface Plasmon Resonance (SPR) using a Biacore T200 from GE Healthcare. N-terminally biotinylated nsp14 (aa 1-527) and C-terminally biotinylated SETD3 (aa 1-605, as control), were coupled on a CM5 SPR Sensor chip (GE healthcare). Compounds were injected into the sensitised chip at 5 concentrations (0.6, 1.9, 5.6, 16.7 and 50 μM) plus 0.5% DMSO at 50 μL/min, using HBS-EP Plus buffer (10 mM HEPES pH 7.4, 150 mM NaCl, 2 mM EDTA, 0.005 % Tween 20), and 0.5 % DMSO. Compounds with lower IC_50_ values (SS148, SAH, WZ16 and JL27-56A1) were injected at the following concentrations: 0.06, 0.19, 0.56, 1.67, and 5 μM. Contact time was 60 seconds, and disassociation time was 120 seconds. Buffer alone (plus 0.5% DMSO) was used for blank injections; and buffers containing 0.4 to 0.6 % DMSO were used for buffer corrections.

### Modeling

DS0464 was docked into the methyltransferase active site of SARS-CoV-1 nsp14 (PDB code 5C8S) with ICM (Molsoft, San Diego). PDB coordinates were loaded, side-chains that were missing due to poor electron-density were built and energy-minimized with a biased-probability Monte Carlo simulation, hydrogens were added and rotameric states of hydroxyl groups, terminal amides and histidine side-chains were optimized in the internal coordinate space. Ligand docking was conducted with fully flexible ligand and a grid representation of the protein with a Monte Carlo-based energy minimization^43^.

## Acknowledgments

We thank Dr. Aled Edwards and Dr. Cheryl Arrowsmith for continued support. This research was funded by the University of Toronto COVID-19 Action Initiative-2020 and COVID-19 Mitacs Accelerate postdoctoral awards to AKY and SP; the US National Institutes of Health grant (R35GM131858) to Dr. Minkui Luo; and NIH/NIAID contract HHSN272201700060C to Dr. Karla Satchell. The Structural Genomics Consortium is a registered charity (no: 1097737) that receives funds from; AbbVie, Bayer Pharma AG, Boehringer Ingelheim, Canada Foundation for Innovation, Eshelman Institute for Innovation, Genentech, Genome Canada through Ontario Genomics Institute [OGI-196], EU/EFPIA/OICR/McGill/KTH, Diamond Innovative Medicines Initiative 2 Joint Undertaking [EUbOPEN grant 875510], Janssen, Merck KGaA (aka EMD in Canada and US), Merck & Co (aka MSD outside Canada and US), Pfizer, São Paulo Research Foundation-FAPESP, Takeda and Wellcome [106169/ZZ14/Z].

## Author contributions

KD designed and performed experiments, analyzed data and wrote the manuscript, M.S performed all computational analysis and wrote the manuscript, S.P, A.K.Y, FL, and IC designed experiments and contributed to assays and screening, PG and TH purified proteins, PL cloned the expression construct, AB prepared compounds and screening materials, K.J.F.S provided reagents, KW, DL, JL, DS, ML, JJ, PB, PVF synthesized compounds and provided compound libraries, M.V was the Principal Investigator, conceived the idea, lead the overall project, designed experiments, reviewed data and wrote the manuscript. All authors reviewed the manuscript.

## Supplementary Data

### Protein Expression and Purification

The expression clone for nsp14 with a 6xHis-tag and tobacco etch virus (TEV) protease cleavage site was codon optimized, synthesized, and cloned into the NdeI-BamH1 sites of pET11b by Bio Basic Inc. (Markham, ON, Canada). A DNA fragment encoding full-length SARS-CoV-2 nsp14 (A1 to Q527) was amplified from this synthetic template by PCR and sub-cloned into p28BIOH-LIC (Addgene plasmid # 62352). This added a N-terminal Avi-tag for in vivo biotinylation and C-terminal His-tag. Following transformation into E. coli BL21 (DE3) containing a BirA expression plasmid (BPS Bioscience) the cells were cultured in LB broth overnight at 37 °C supplemented with kanamycin and spectinomycin. Terrific Broth was inoculated with overnight culture of nsp14 supplemented with 50 μg/mL Kanamycin and 100 μg/mL spectinomycin and the cells were grown using LEX system (https://www.thesgc.org/science/lex). At OD600 of 0.8-1.5, the temperature was lowered to 18°C and the culture was supplemented with 1 mM IPTG (isopropyl-1-thio-D-galactopyranoside). D-Biotin was added at 10 μg/mL final concentration to allow in cell biotinylation of the protein. The cells were incubated overnight before being harvested (5000 ×g for 15 minutes at 4°C) using a Beckman Coulter centrifuge (model: Avanti J-20 X PI, rotor JLA-8.1000).

Harvested cells were re-suspended in binding buffer containing 20 mM Tris-HCl pH 8.5, 500 mM NaCl, 10 mM MgCl2, 5% glycerol, and 0.5 mM TCEP. The mixture was then treated with 1x protease inhibitor cocktail (100X protease inhibitor cocktail in 70% ethanol (0.25 mg/ml Aprotinin, 0.25 mg/ml Leupeptin, 0.25 mg/ml Pepstatin A and 0.25 mg/ml E-64)) and Roche complete EDTA-free protease inhibitor cocktail tablets. The cells were lysed chemically by rotating 30 min with CHAPS (final concentration of 0.5%) and 5 μl/L Benzonase Nuclease (in house) followed by sonication at frequency of 8.0 (10” on/7” off) for 10 min (Sonicator 3000, Misoni). The crude extract was clarified by high-speed centrifugation (45 min at 36,000 ×g at 4°C) by Beckman Coulter centrifuge (rotor: JLA 16.250).

The clarified lysate was passed through pre-equilibrated Monomeric Avidin agarose resin (Pierce) following the manufacturer’s instructions. Briefly, the column was washed with PBS followed by blocking the resin with 2 mM Biotin solution in PBS. Subsequently, the excess biotin was removed by 0.1 M Glycine, pH 2.8 followed by a full column wash with PBS. The lysate was continuously passed through the beads for 3 hours at 4°C. The resin was then washed with PBS to remove the non-specific proteins until Bradford dye remained unchanged. The biotinylated nsp14 was then eluted using 2 mM biotin solution in PBS. Finally, the purity of the nsp14 protein was confirmed by SDS-PAGE (**Supplementary Figure 17a**).

The eluted protein was then dialyzed into the final buffer containing 50 mM Tris-HCl pH 8.5, 200 mM NaCl, 5% glycerol, 0.5 mM TCEP for 60 min while spinning at 4°C.

To evaluate the stability of nsp14 protein after freeze and thaw, an analytical SEC was performed by first thawing the protein on ice and centrifuging it at 18,900 ×g for 10 min at 4°C. Then, the nsp14 was loaded onto Superdex200 10X 300 column (GE Healthcare) after equilibration with 50 mM Tris-HCl pH 8.0 buffer containing 200 mM NaCl, 5% glycerol, 0.5 mM TCEP and the molecular weight of the protein was estimated based on the standard peaks used to plot the calibration curve. The molecular weight of the nsp14 was confirmed by running 10 μg of proteins on mass spectrometer (Agilent Technologies, 6545 Q-TOF LC/MS) (**Supplementary Figure 17b**).

**Supplementary Figure 1:**
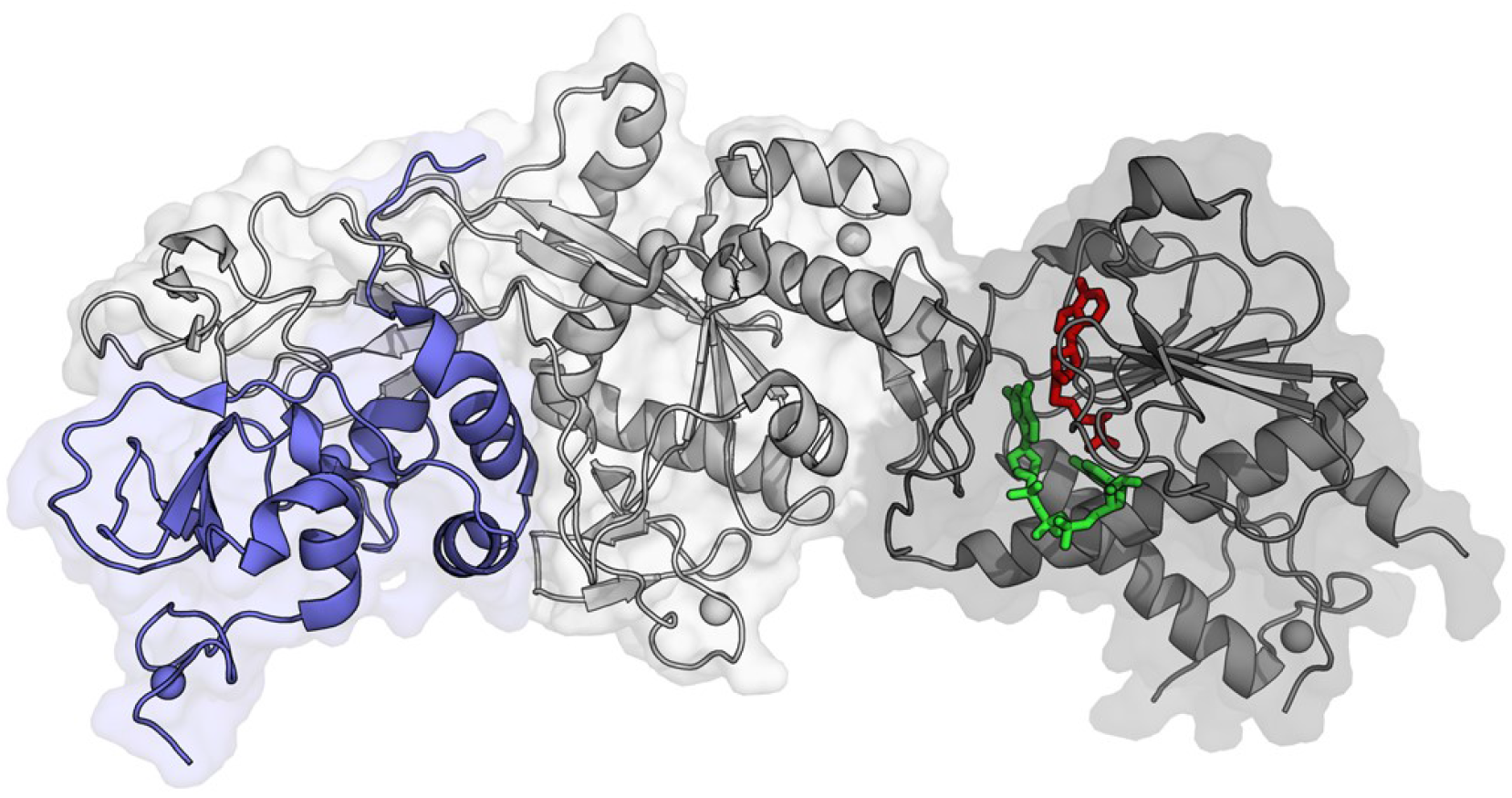
Crystal structure of SARS-CoV nsp10-nsp14 in complex with GpppA and SAH. Exonuclease domain of nsp14 is in light gray, methyltransferase domain of nsp14 is in dark gray, SAH in red stick, and GpppA with green stick. Nsp10 is shown in blue. (PDB ID: 5C8S)^1^

**Supplementary Figure 2:**
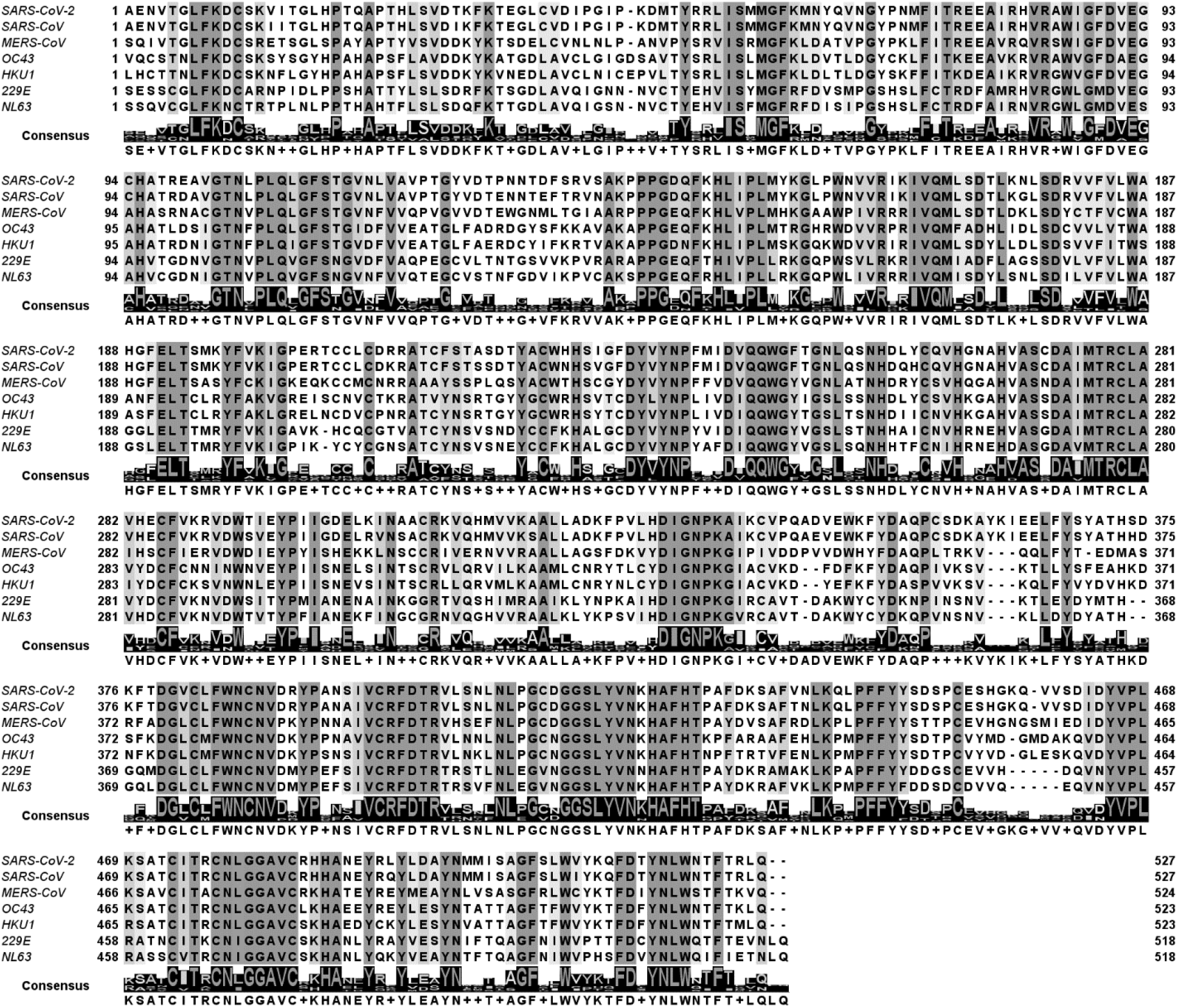
Sequence alignment of nsp14 from 7 coronaviruses that are currently known to infect humans. The amino acid sequence of nsp14 is highly conserved among these coronaviruses. The alignment indicates 95.1, 62.7, 57.8, 58.5, 52.9 and 53.7% identity of SARS-CoV-2 nsp14 sequence with SARS-CoV, MERS-CoV, OC43, HKU1, 299E and NL63, respectively. Total of 184 amino acids are conserved among all 7 coronaviruses. The sequences have been aligned using Clustal Omega.^2^ The image was prepared using Jalview.

**Supplementary Figure 3:**
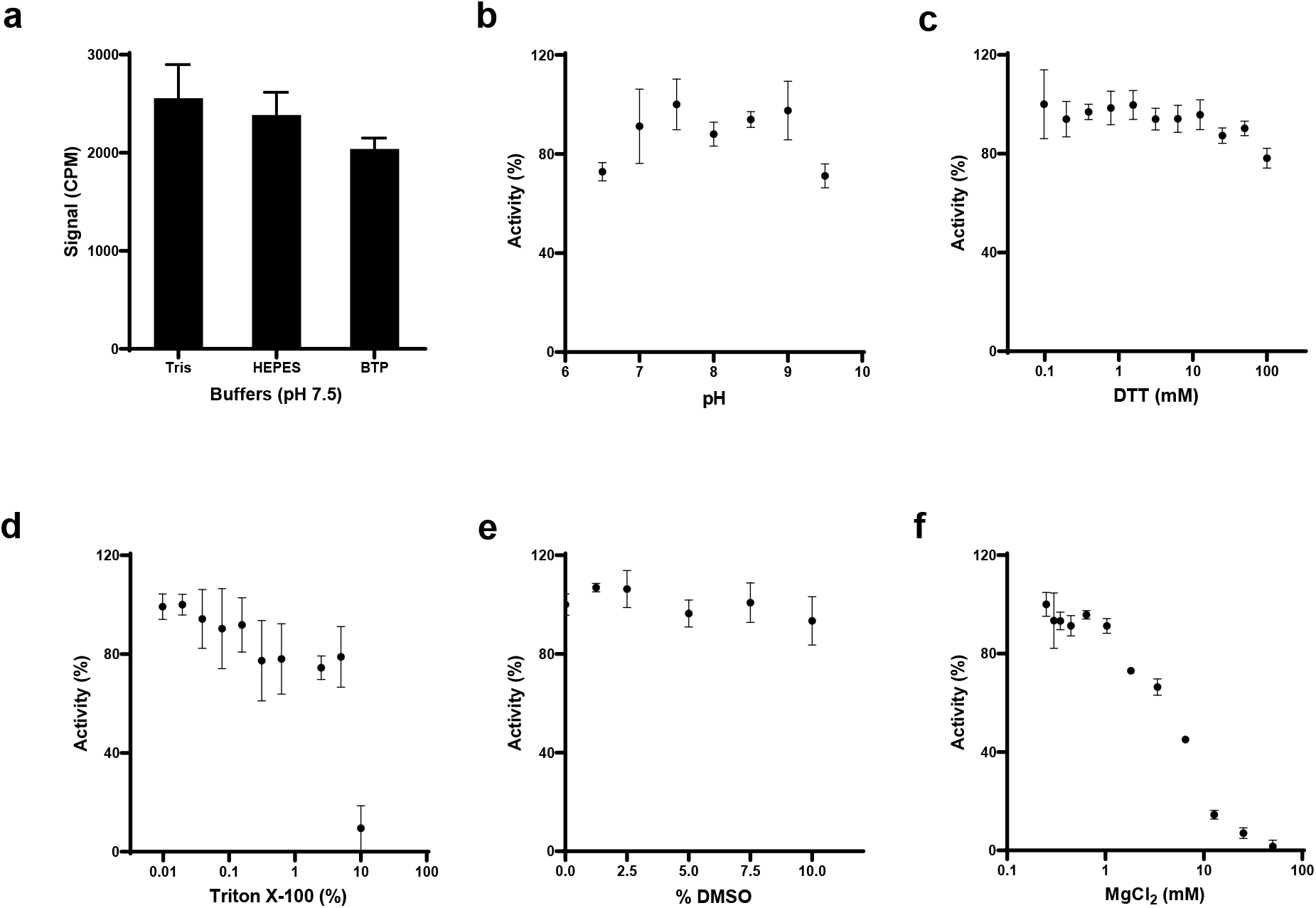
Nsp14 activity assay optimization. Effect of (a) Buffers, (b) pH, (c) DTT, (d) Triton X-100, (e) DMSO and (f) MgCl2 on nsp14 activity was assessed. All values are mean ± standard deviation of three independent experiments (n=3).

**Supplementary Figure 4:**
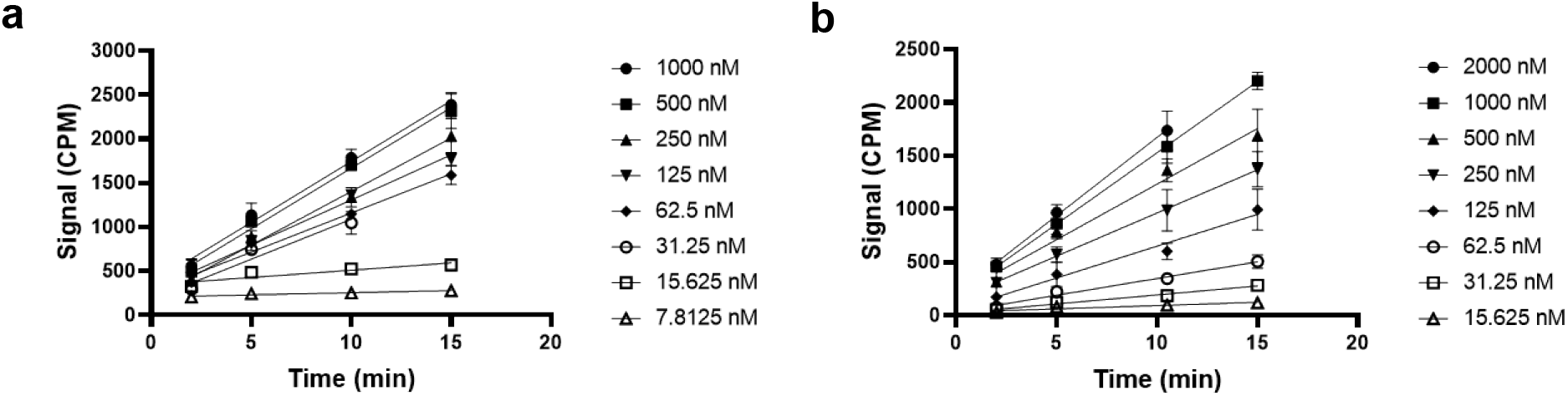
Kinetic characterization of nsp14 activity. Linear initial velocities at various concentrations of (a) RNA substrate at fixed concentration of SAM (1 μM) and (b) various concentrations of SAM at fixed concentration of RNA (1 μM) were evaluated. The values calculated within the linear period for each reaction were used to plot figure 1a and 1b and calculate the K_m_ and k_cat_ values. All values are mean ± standard deviation of three independent experiments (n=3).

**Supplementary Figure 5:**
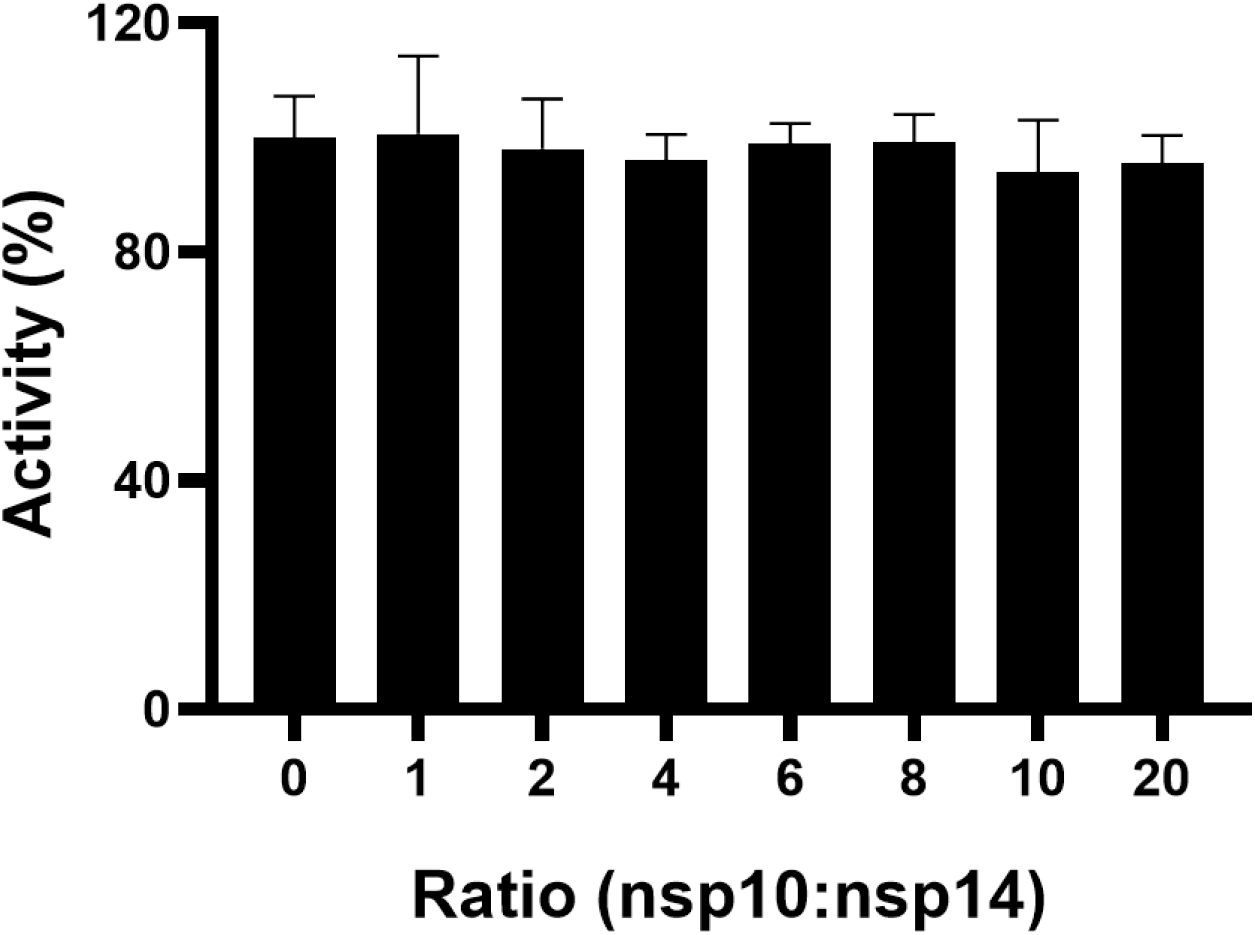
Effect of nsp10-nsp14 complex formation on methyltransferase activity of nsp14. Complex preparations of nsp10: nsp14 ratios up to 20 (nsp10): 1 (nsp14) were tested for methyltransferase activity. No significant effect on activity of nsp14 was observed at any concentrations of nsp10.

**Supplementary Figure 6:**
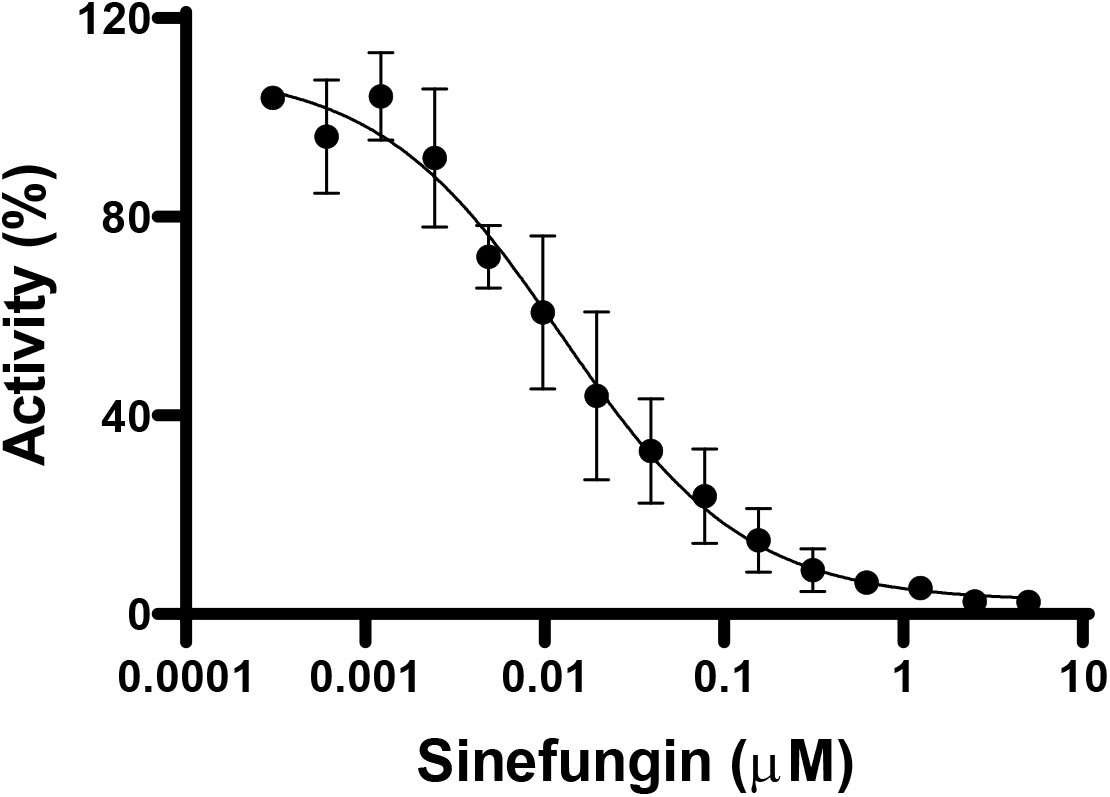
Dose response for nsp14 inhibition by sinefungin. All values are mean ± standard deviation of three independent experiments (n=3). IC_50_ value (0.019 ± 0.01 μM) was determined for sinefungin at optimized conditions as described in methods.

**Supplementary Figure 7:**
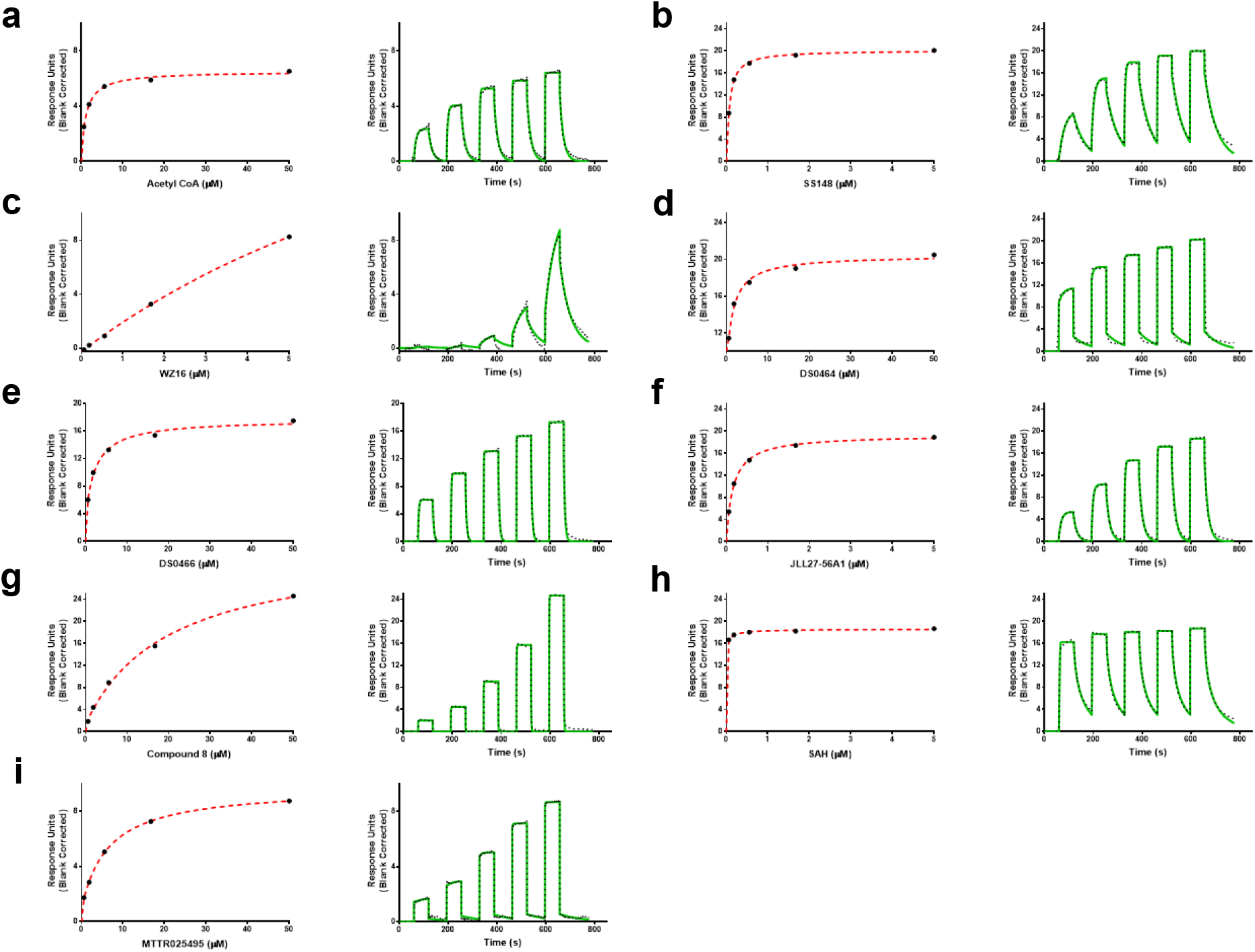
Orthogonal confirmation of nsp14 screening hits. The screening hits, (a) Acetyl CoA, (b) SS148, (c) WZ16, (d) DS0464, (e) DS0466, (f) JL27-56A1, (g) Compound 8, (h) SAH and (i) MTTR025495 were tested by Surface Plasmon Resonance (SPR). For each compound, the Sensorgram (solid green) is shown with the kinetic fit (black dots), and the steady state response (black circles) with the steady state 1:1 binding model fitting (red dashed line). K_D_ values are presented in Table 1.

**Supplementary Figure 8:**
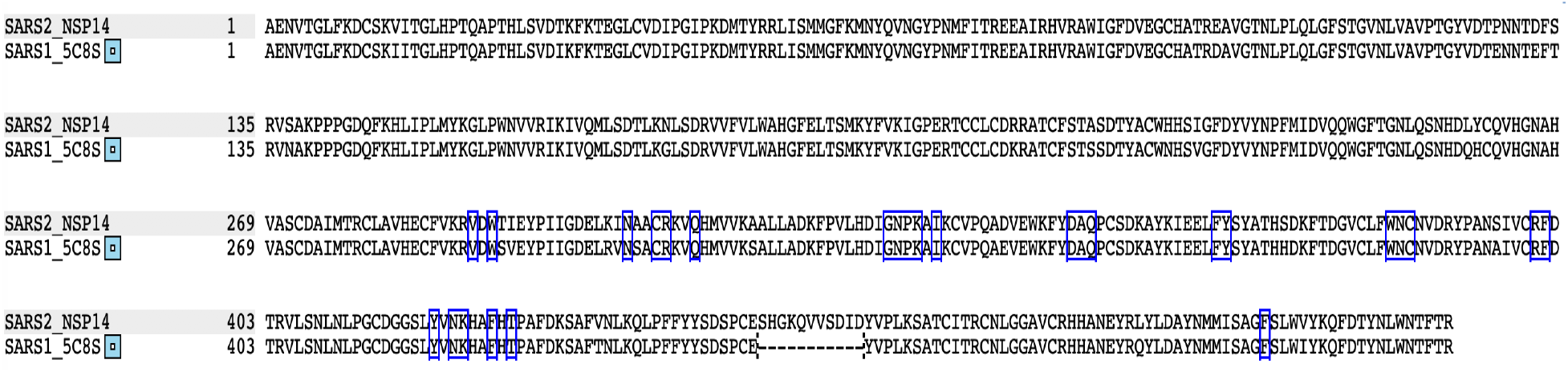
Amino acid homology of nsp14 from SARS-CoV-1 and SARS-CoV-2. Sequence alignment of SARS-CoV-1 nsp14 from PDB code 5C8S^1^ and SARS-CoV-2 nsp14 downloaded from Uniprot. Amino acids within 4.0 Å of the cofactor or GpppA in structure 5C8S are boxed in blue.

**Supplementary Figure 9:**
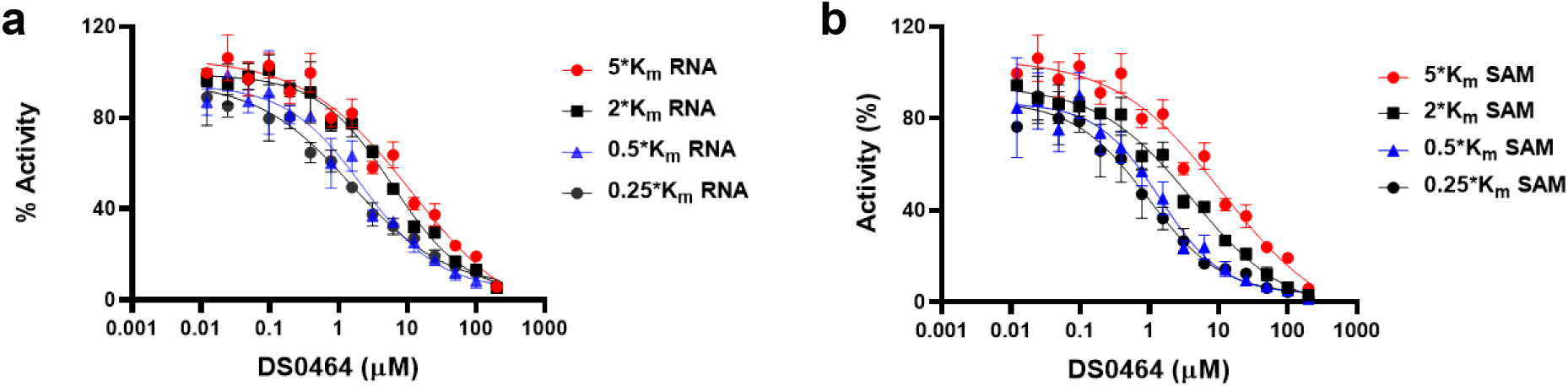
MOA determination. Dose Response Curves for DS0464 against nsp14 at varying concentrations of (a) RNA at 1.25 μM of SAM and (b) SAM at 250 nM of RNA (saturating concentration). The IC_50_ values from these experiments were plotted in figure 5 for determining the mechanism of action of DS0464.

**Supplementary Figure 10:**
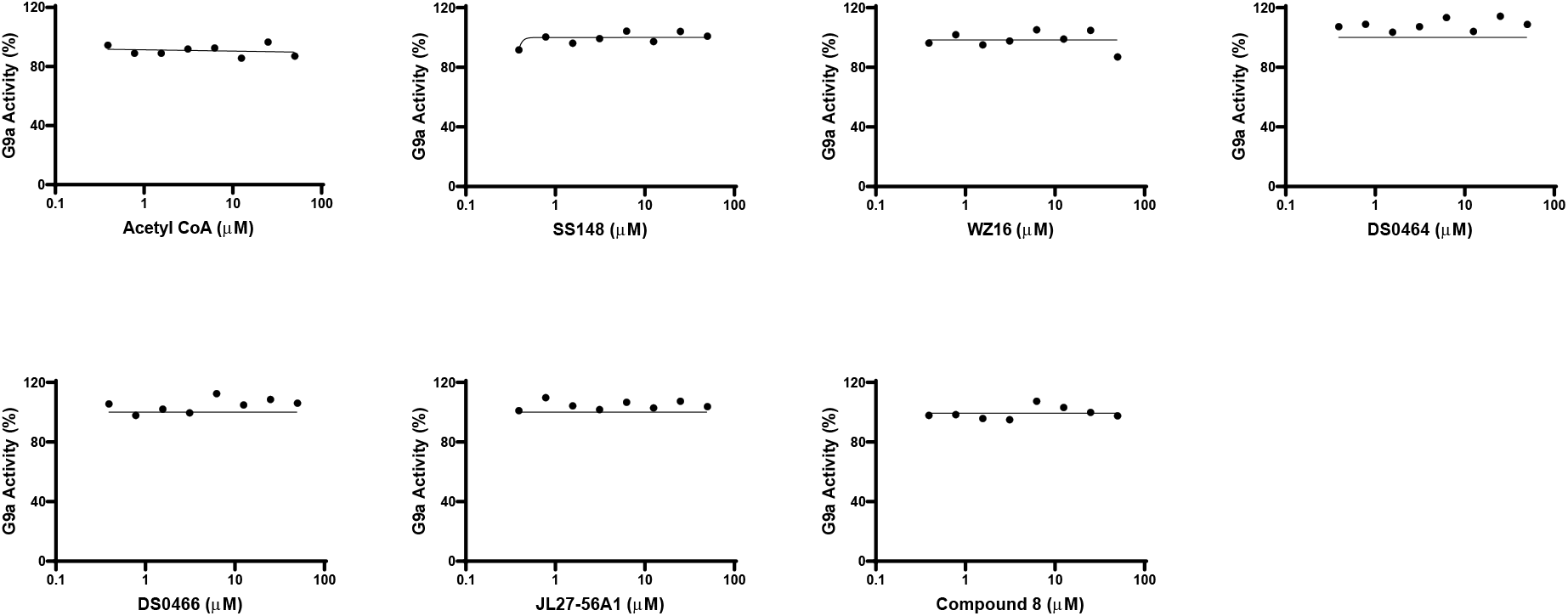
Selectivity of the nsp14 screening hits against G9a, a protein lysine methyltransferase (PKMT) . Dose response data are presented for indicated compounds. None of the compounds inhibit G9a activity up to 50 μM of compound.

**Supplementary Figure 11:**
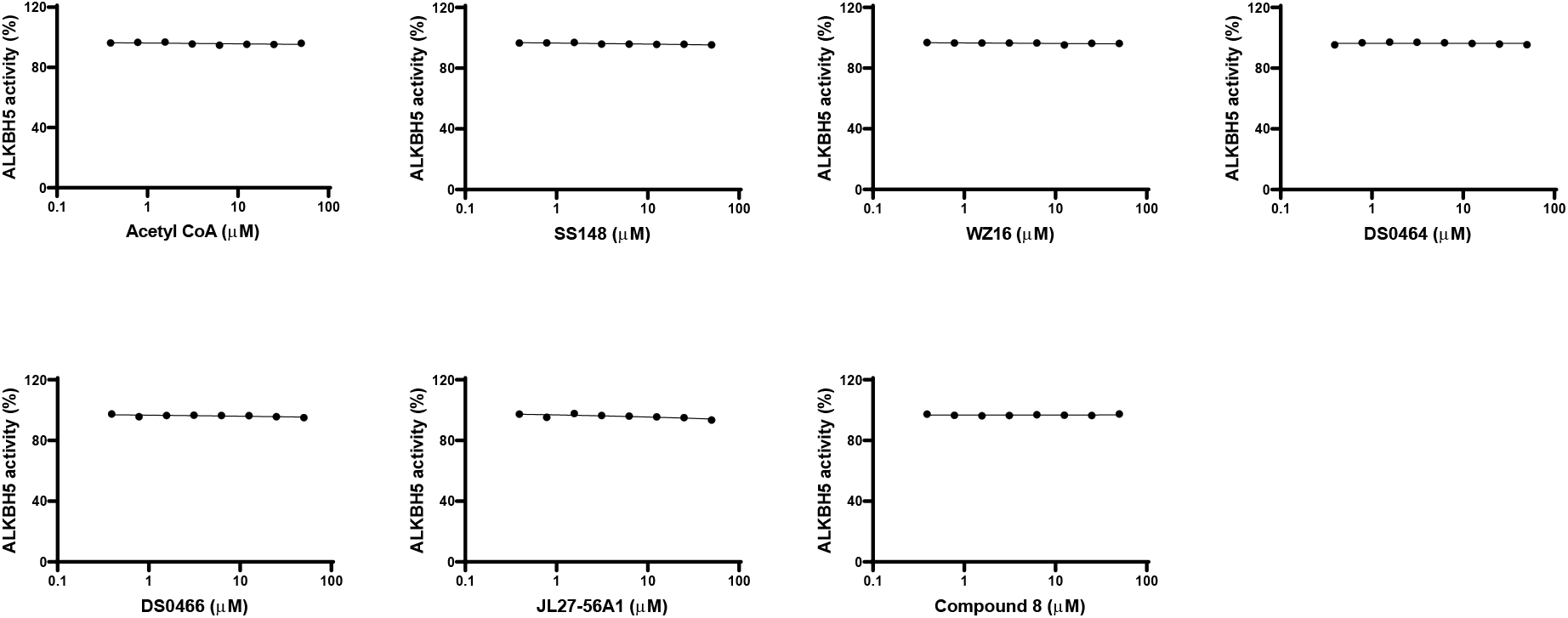
Selectivity of the nsp14 screening hits against ALKBH5, an RNA demethylase. Dose response data are presented for indicated compounds. None of the compounds inhibit ALKBH5 activity up to 50 μM of compound.

**Supplementary Figure 12:**
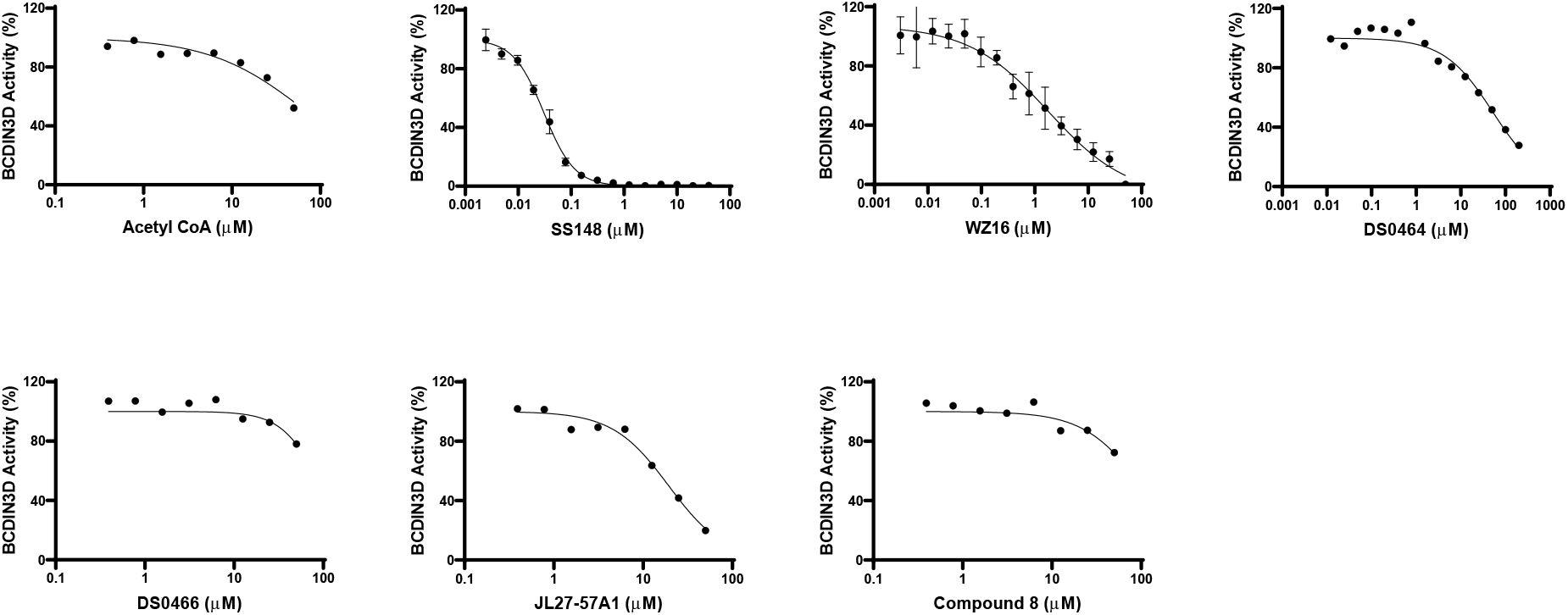
Selectivity of the nsp14 screening hits against BCDIN3D, an RNA methyltransferase (RNMT). Dose response data are presented for indicated compounds. Experiments were performed in triplicate only for SS148 and WZ16.

**Supplementary Figure 13:**
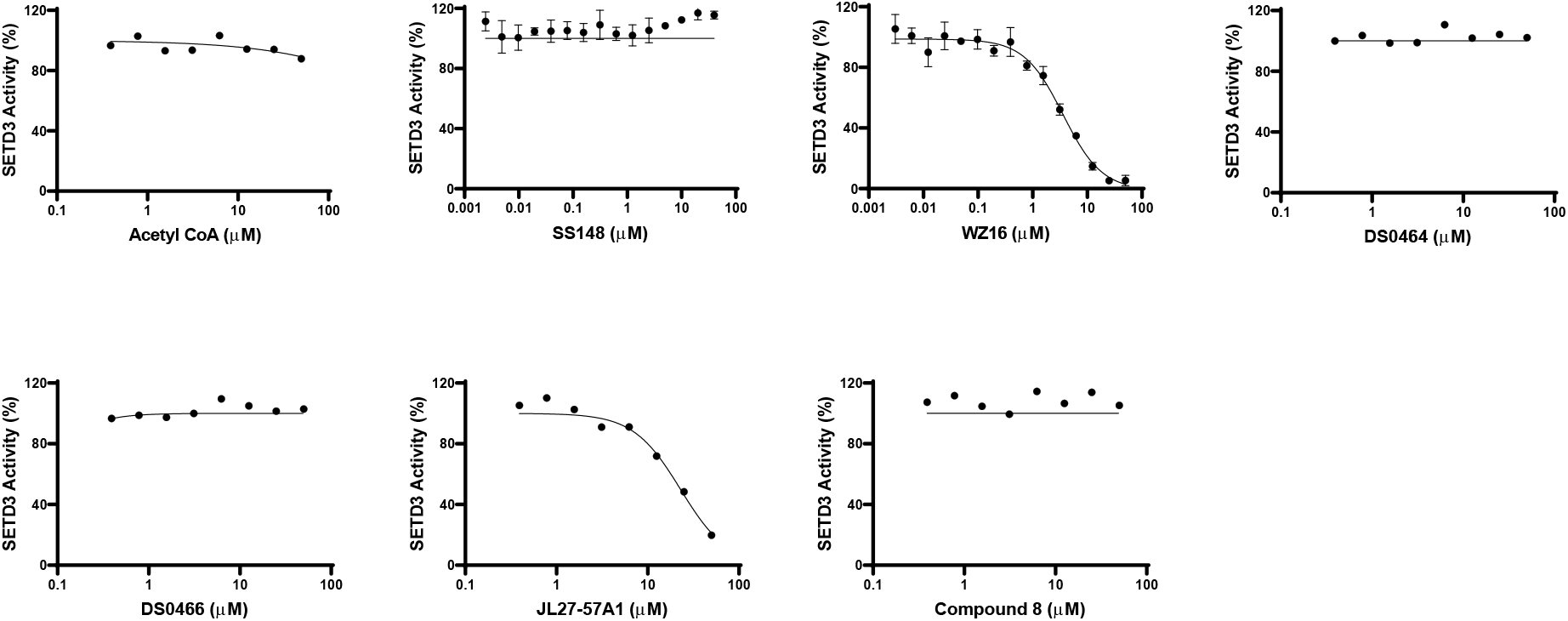
Selectivity of the nsp14 screening hits against SETD3, a PKMT. Dose response data are presented for indicated compounds. Experiments were performed in triplicate for SS148 and WZ16 (n=3).

**Supplementary Figure 14:**
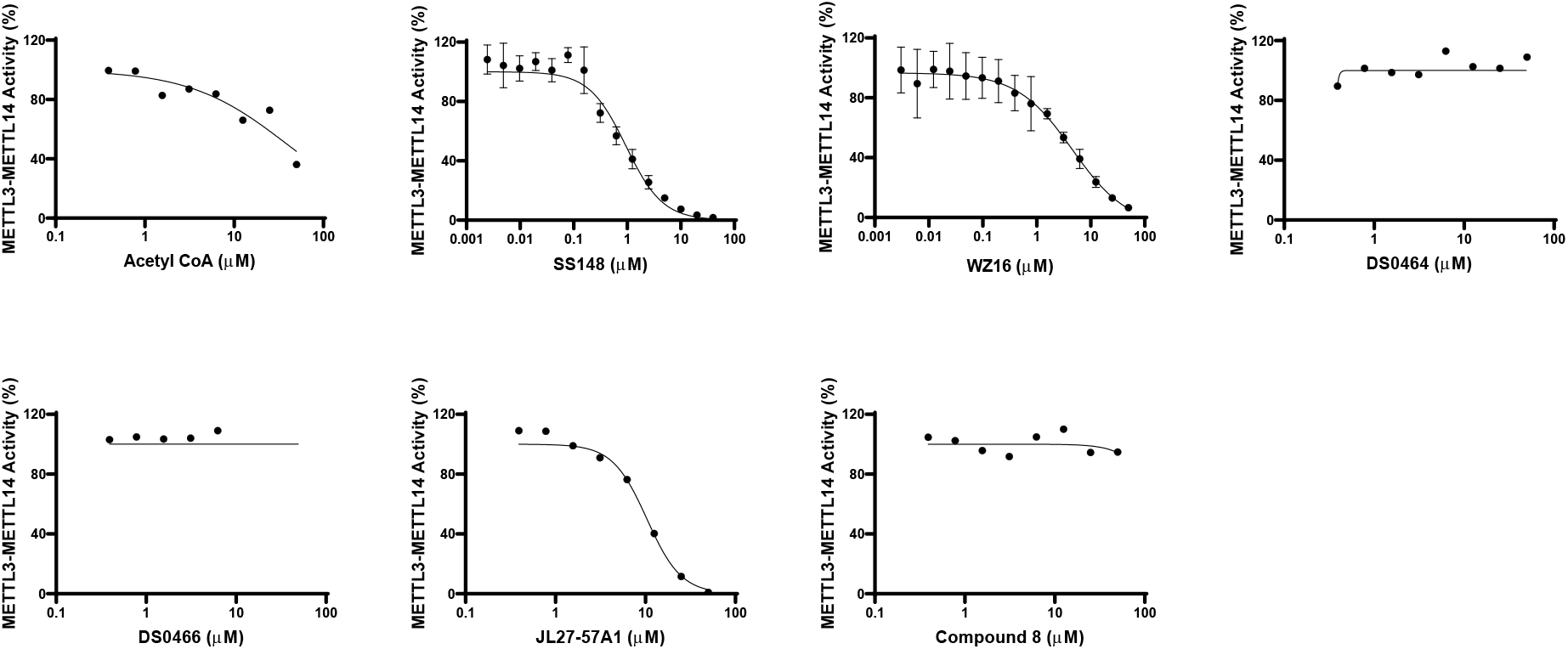
Selectivity of the nsp14 screening hits against METTL3-METTL14, an RNA methyltransferase. Dose response data are presented for nsp14 screening hits. Experiments were performed in triplicate for SS148 and WZ16 (n=3).

**Supplementary Figure 15:**
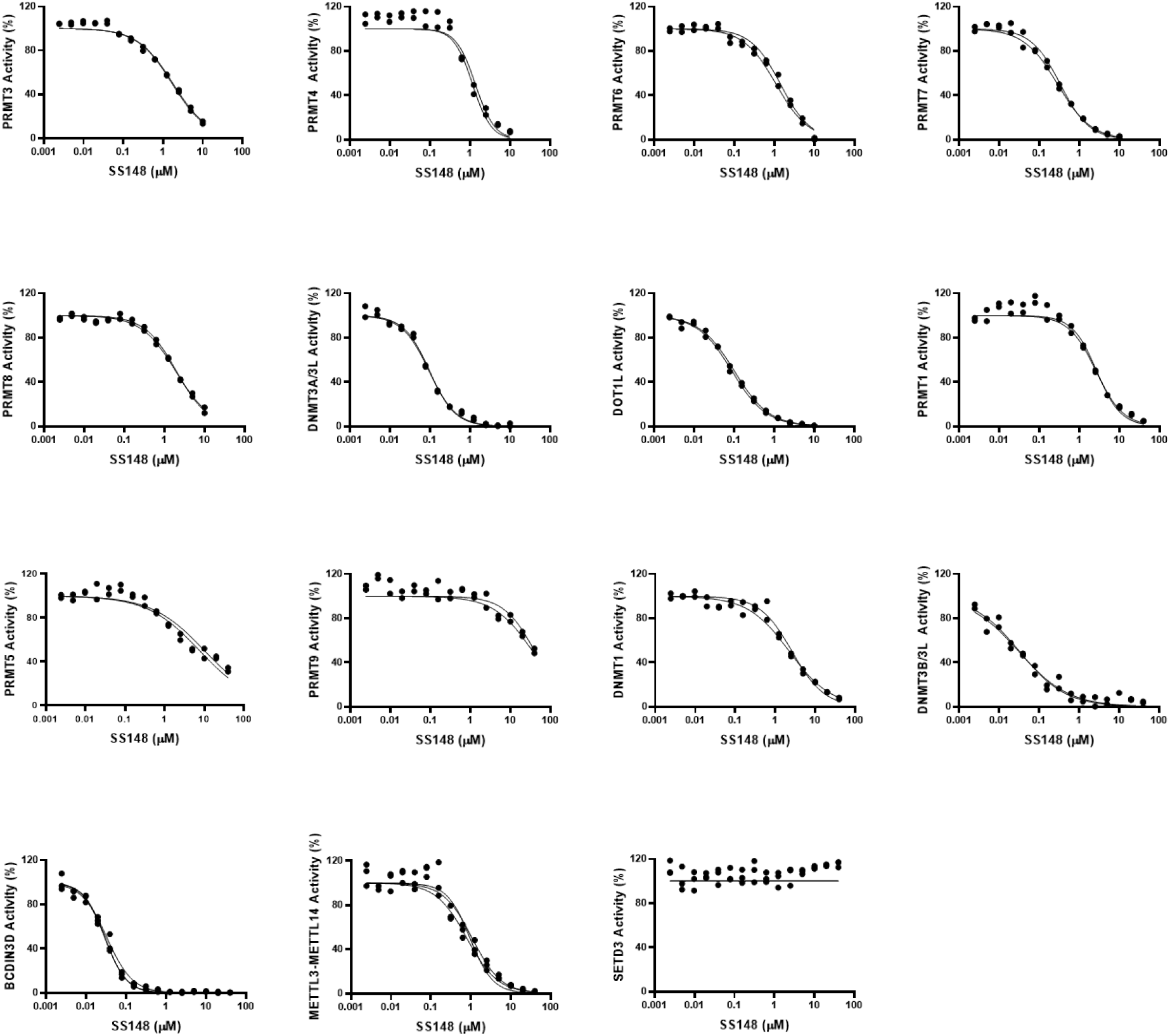
Selectivity of SS148. Selectivity of SS148 against protein arginine methyltransferases (PRMT1, PRMT3, PRMT4, PRMT5, PRMT6, PRMT7, PRMT8, and PRMT9), Protein Lysine methyltransferase (DOT1L and SETD3), DNA methyltransferases (DNMT1, DNMT3a, and DNMT3b) and RNA methyltransferase (BCDIN3D and METTL3-METTL14) were assessed in dose response. Experiments were performed in duplicate (n=2).

**Supplementary Figure 16:**
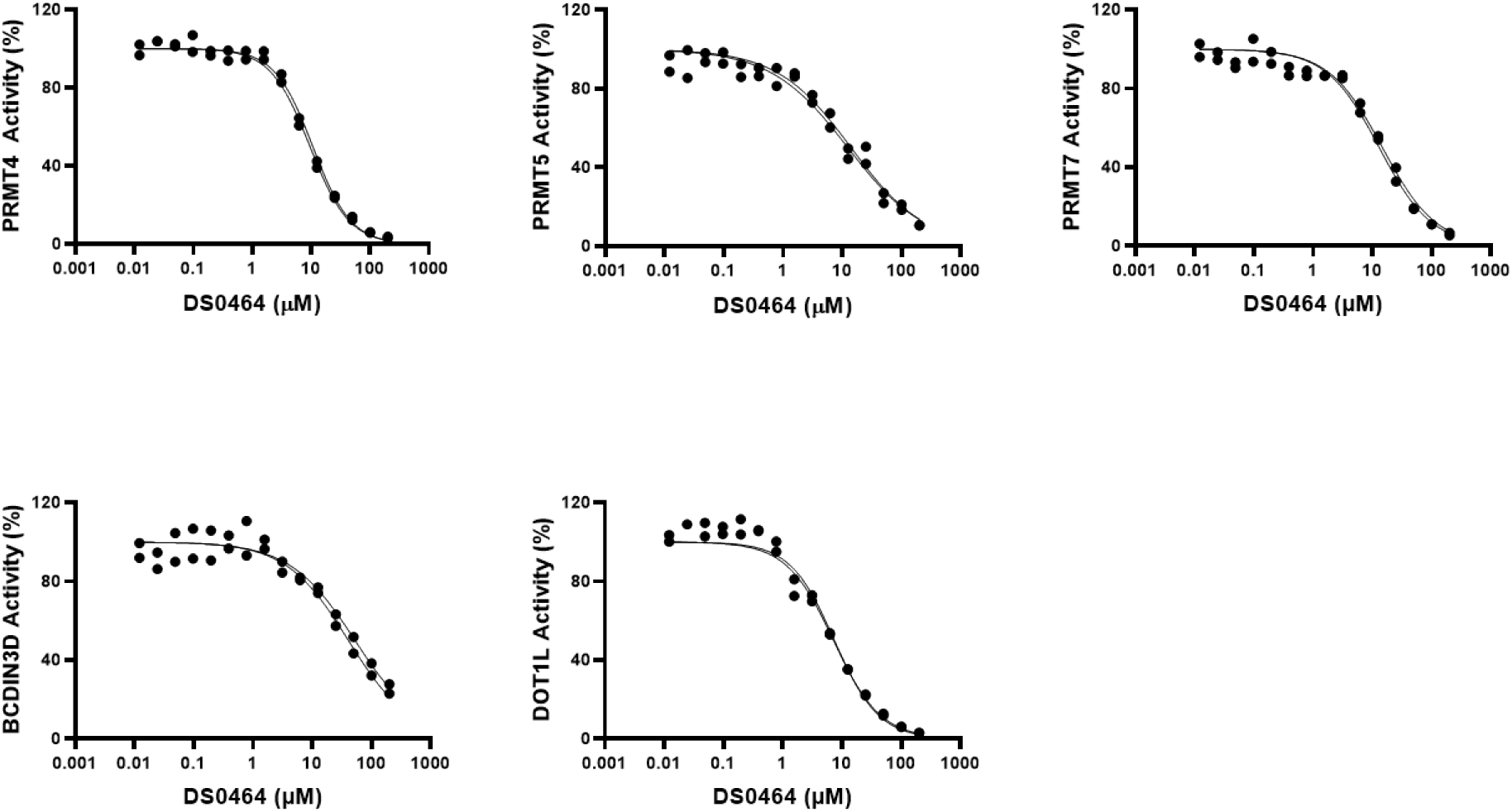
Selectivity of DS0464. Dose response analysis of DS0464 against selected methyltransferases. Experiments were performed in duplicate (n=2).

**Supplementary Figure 17:**
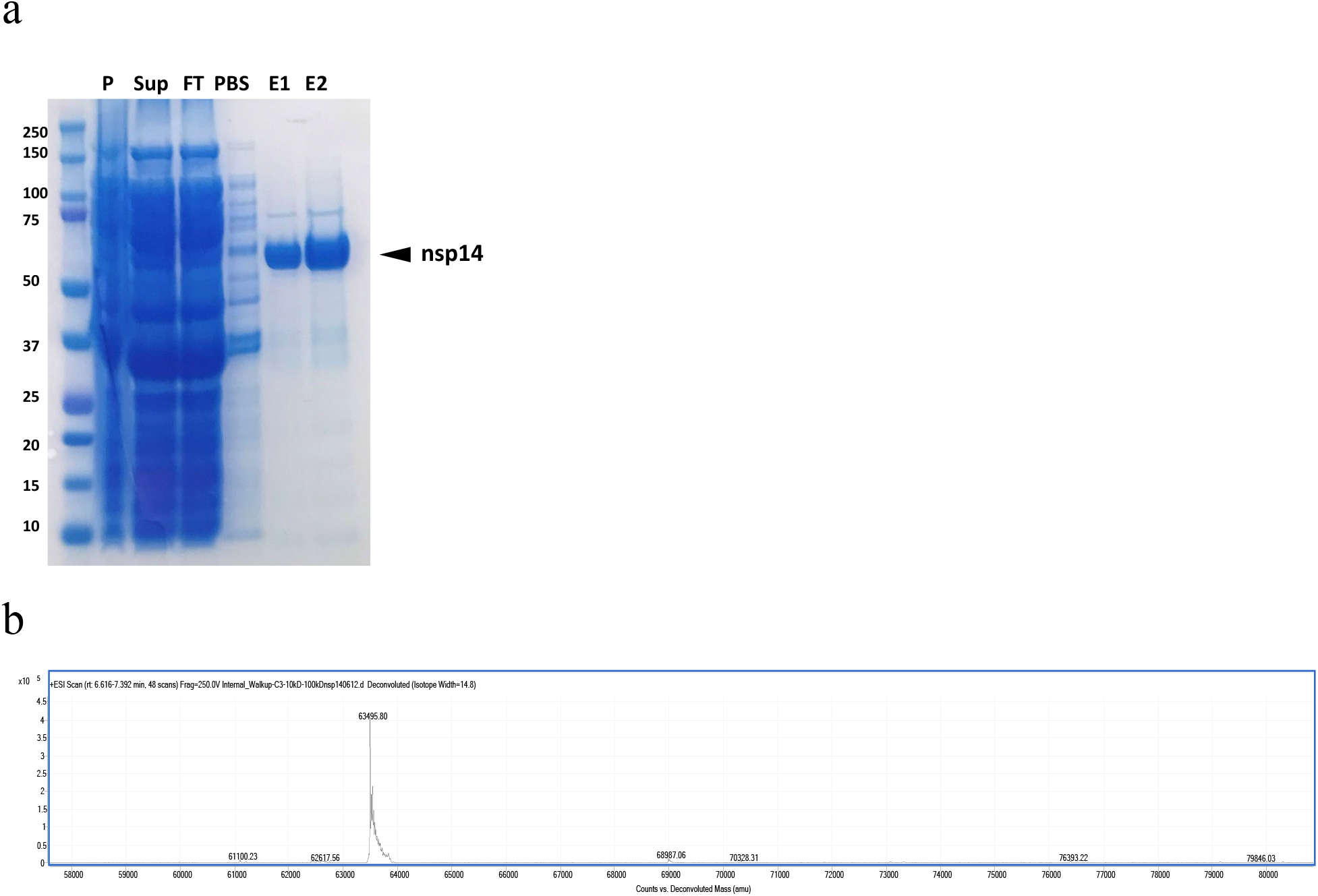
Nsp14 purification and quality assessment. (**a**) SDS-PAGE demonstrating biotinylated nsp14 after affinity purification on Avidin agarose resin. The lanes next to the molecular-weight size marker from left to right correspond to (1) pellet, (2) supernatant, (3) flow-through, (4) wash, (5) 10 μL elution, and (6) 15 μL elution. The arrowhead indicates the nsp14 at approximately 63 kDa. (**b**) Mass spectrum of nsp14 confirms the molecular weight of the protein.

## References

1. Zhu, N. et al. A Novel Coronavirus from Patients with Pneumonia in China, 2019. N Engl J Med 382, 727–733 (2020).

2. Adams, M.J. et al. Ratification vote on taxonomic proposals to the International Committee on Taxonomy of Viruses (2016). Arch Virol 161, 2921–49 (2016).

3. Coronaviridae Study Group of the International Committee on Taxonomy of, V. The species Severe acute respiratory syndrome-related coronavirus: classifying 2019-nCoV and naming it SARS-CoV-2. Nat Microbiol 5, 536–544 (2020).

4. Chan-Yeung, M. & Xu, R.H. SARS: epidemiology. Respirology 8 Suppl, S9–14 (2003).

5. Zaki, A.M., van Boheemen, S., Bestebroer, T.M., Osterhaus, A.D. & Fouchier, R.A. Isolation of a novel coronavirus from a man with pneumonia in Saudi Arabia. N Engl J Med 367, 1814–20 (2012).

6. Chan, J.F. et al. Genomic characterization of the 2019 novel human-pathogenic coronavirus isolated from a patient with atypical pneumonia after visiting Wuhan. Emerg Microbes Infect 9, 221–236 (2020).

7. van Boheemen, S. et al. Genomic characterization of a newly discovered coronavirus associated with acute respiratory distress syndrome in humans. mBio 3, e00473–12 (2012).

8. Chen, Y., Liu, Q. & Guo, D. Emerging coronaviruses: Genome structure, replication, and pathogenesis. J Med Virol 92, 418–423 (2020).

9. Baez-Santos, Y.M., St John, S.E. & Mesecar, A.D. The SARS-coronavirus papain-like protease: structure, function and inhibition by designed antiviral compounds. Antiviral Res 115, 21–38 (2015).

10. Chen, Y. & Guo, D. Molecular mechanisms of coronavirus RNA capping and methylation. Virol Sin 31, 3–11 (2016).

11. Eckerle, L.D. et al. Infidelity of SARS-CoV Nsp14-exonuclease mutant virus replication is revealed by complete genome sequencing. PLoS Pathog 6, e1000896 (2010).

12. Bouvet, M. et al. RNA 3’-end mismatch excision by the severe acute respiratory syndrome coronavirus nonstructural protein nsp10/nsp14 exoribonuclease complex. Proc Natl Acad Sci U S A 109, 9372–7 (2012).

13. Smith, E.C. et al. Mutations in coronavirus nonstructural protein 10 decrease virus replication fidelity. J Virol 89, 6418–26 (2015).

14. Subissi, L. et al. One severe acute respiratory syndrome coronavirus protein complex integrates processive RNA polymerase and exonuclease activities. Proc Natl Acad Sci U S A 111, E3900–9 (2014).

15. Decroly, E. et al. Crystal structure and functional analysis of the SARS-coronavirus RNA cap 2’-O-methyltransferase nsp10/nsp16 complex. PLoS Pathog 7, e1002059 (2011).

16. Bouvet, M. et al. In vitro reconstitution of SARS-coronavirus mRNA cap methylation. PLoS Pathog 6, e1000863 (2010).

17. Chen, Y. et al. Functional screen reveals SARS coronavirus nonstructural protein nsp14 as a novel cap N7 methyltransferase. Proc Natl Acad Sci U S A 106, 3484–9 (2009).

18. Marcotrigiano, J., Gingras, A.C., Sonenberg, N. & Burley, S.K. Cocrystal structure of the messenger RNA 5’ cap-binding protein (eIF4E) bound to 7-methyl-GDP. Cell 89, 951–61 (1997).

19. Decroly, E., Ferron, F., Lescar, J. & Canard, B. Conventional and unconventional mechanisms for capping viral mRNA. Nat Rev Microbiol 10, 51–65 (2011).

20. Ivanov, K.A. & Ziebuhr, J. Human coronavirus 229E nonstructural protein 13: characterization of duplex-unwinding, nucleoside triphosphatase, and RNA 5’-triphosphatase activities. J Virol 78, 7833–8 (2004).

21. Ivanov, K.A. et al. Multiple enzymatic activities associated with severe acute respiratory syndrome coronavirus helicase. J Virol 78, 5619–32 (2004).

22. Decroly, E. et al. Coronavirus nonstructural protein 16 is a cap-0 binding enzyme possessing (nucleoside-2’O)-methyltransferase activity. J Virol 82, 8071–84 (2008).

23. Corman, V.M., Muth, D., Niemeyer, D. & Drosten, C. Hosts and Sources of Endemic Human Coronaviruses. Adv Virus Res 100, 163–188 (2018).

24. Scheer, S. et al. A chemical biology toolbox to study protein methyltransferases and epigenetic signaling. Nat Commun 10, 19 (2019).

25. Campagna-Slater, V. et al. Structural chemistry of the histone methyltransferases cofactor binding site. J Chem Inf Model 51, 612–23 (2011).

26. Ferron, F., Decroly, E., Selisko, B. & Canard, B. The viral RNA capping machinery as a target for antiviral drugs. Antiviral Res 96, 21–31 (2012).

27. Ferreira de Freitas, R., Ivanochko, D. & Schapira, M. Methyltransferase Inhibitors: Competing with, or Exploiting the Bound Cofactor. Molecules 24, 10.3390/molecules24244492 (2019).

28. Stein, E.M. et al. The DOT1L inhibitor pinometostat reduces H3K79 methylation and has modest clinical activity in adult acute leukemia. Blood 131, 2661–2669 (2018).

29. Gounder, M. et al. Tazemetostat in advanced epithelioid sarcoma with loss of INI1/SMARCB1: an international, open-label, phase 2 basket study. Lancet Oncol 21, 1423–1432 (2020).

30. Rohman, M. & Wingfield, J. High-Throughput Screening Using Mass Spectrometry within Drug Discovery. Methods Mol Biol 1439, 47–63 (2016).

31. Haslam, C. et al. The Evolution of MALDI-TOF Mass Spectrometry toward Ultra-High-Throughput Screening: 1536-Well Format and Beyond. J Biomol Screen 21, 176–86 (2016).

32. Winter, M. et al. Automated MALDI Target Preparation Concept: Providing Ultra-High-Throughput Mass Spectrometry-Based Screening for Drug Discovery. SLAS Technol 24, 209–221 (2019).

33. Janzen, W.P. Screening technologies for small molecule discovery: the state of the art. Chem Biol 21, 1162–70 (2014).

34. Yu, W. et al. Catalytic site remodelling of the DOT1L methyltransferase by selective inhibitors. Nat Commun 3, 1288 (2012).

35. Daigle, S.R. et al. Selective killing of mixed lineage leukemia cells by a potent small-molecule DOT1L inhibitor. Cancer Cell 20, 53–65 (2011).

36. Konze, K.D. et al. An orally bioavailable chemical probe of the Lysine Methyltransferases EZH2 and EZH1. ACS Chem Biol 8, 1324–34 (2013).

37. Taylor, A.P. et al. Selective, Small-Molecule Co-Factor Binding Site Inhibition of a Su(var)3-9, Enhancer of Zeste, Trithorax Domain Containing Lysine Methyltransferase. J Med Chem 62, 7669–7683 (2019).

38. Spurr, S.S. et al. New small molecule inhibitors of histone methyl transferase DOT1L with a nitrile as a non-traditional replacement for heavy halogen atoms. Bioorg Med Chem Lett 26, 4518–4522 (2016).

39. Cai, X.C. et al. A chemical probe of CARM1 alters epigenetic plasticity against breast cancer cell invasion. Elife 8, 8:e47110 (2019).

40. Babault, N. et al. Discovery of Bisubstrate Inhibitors of Nicotinamide N-Methyltransferase (NNMT). J Med Chem 61, 1541–1551 (2018).

41. Hong, S. et al. Nicotinamide N-methyltransferase regulates hepatic nutrient metabolism through Sirt1 protein stabilization. Nat Med 21, 887–94 (2015).

42. Zhang, J.H., Chung, T.D. & Oldenburg, K.R. A Simple Statistical Parameter for Use in Evaluation and Validation of High Throughput Screening Assays. J Biomol Screen 4, 67–73 (1999).

43. Neves, M.A., Totrov, M. & Abagyan, R. Docking and scoring with ICM: the benchmarking results and strategies for improvement. J Comput Aided Mol Des 26, 675–86 (2012).

## REFERENCE LIST

1. Ma, Y. et al. Structural basis and functional analysis of the SARS coronavirus nsp14-nsp10 complex. Proc Natl Acad Sci U S A 112, 9436–41 (2015).

2. Sievers, F. et al. Fast, scalable generation of high-quality protein multiple sequence alignments using Clustal Omega. Mol Syst Biol 7, 539 (2011).

